# Host cyclophilin-mediated maturation of an obligate intracellular bacterial surface virulence factor

**DOI:** 10.1101/2025.10.28.684717

**Authors:** Brandon Sit, Allen G. Sanderlin, Alejandro A. Guzman, Lauren E. Bird, Clara Y. Zhu, Patrick J. Woida, Paige N. McCaslin, Julia Kevil-Yeager, Paige E. Allen, Jason A. Carlyon, Mary M. Weber, Thomas P. Burke, John G. Doench, Rebecca L. Lamason

**Affiliations:** Department of Biology, Massachusetts Institute of Technology, Cambridge, MA, USA; Department of Microbiology and Molecular Genetics, University of California Irvine School of Medicine, Irvine, CA, USA; Department of Microbiology and Immunology, University of Iowa Carver College of Medicine, Iowa City, IA, USA; Department of Microbiology and Immunology, Virginia Commonwealth University Medical Center, School of Medicine, Richmond, VA, USA; Genetic Perturbation Platform, Broad Institute, Cambridge, MA, USA

## Abstract

Adaptation of obligate intracellular bacteria to the eukaryotic cytoplasm creates dependencies on host factors that comprise unique infection mechanisms and vulnerabilities compared to facultative intracellular species. Here, we identify the peptidylprolyl isomerase cyclophilin A (PPIA/CypA) as a convergent host dependency in obligate intracellular bacterial pathogens. Using a host-directed functional genetic screen in the emerging tick-borne pathogen *Rickettsia parkeri*, we find that PPIA promotes infection and actin-based motility through an unprecedented inter-kingdom surface protein maturation event that facilitates transmembrane translocation of the actin-nucleating virulence factor Sca2. PPIA is required for efficient *R. parkeri* infection across human and tick cells, and pharmacological inhibition of PPIA phenocopies the loss of Sca2 *in vitro* and *in vivo*. This dependency extends to *Chlamydia trachomatis* and *Orientia tsutsugamushi* but appears absent in multiple facultative intracellular species. Targeting of host PPIA may thus represent a shared infection dependency across obligate intracellular bacterial pathogens with unmet therapeutic needs.

## Introduction

Intracellular bacterial pathogens rely on numerous interactions with host factors to facilitate infection. These dependencies are particularly important in obligate intracellular bacterial pathogens, a group of evolutionarily diverse microbes that lack the ability to replicate outside of host cells, leaving them entirely reliant on successful host manipulation for survival^1^. The simultaneous loss of factors needed to thrive extracellularly and adaptation to host cytoplasmic niches in obligate intracellular bacteria has driven evolution of remarkable host-dependent infection mechanisms and bacterial physiology^1,2^. These dependencies, which may be useful host-directed therapeutic targets, can be pathogen-specific because of their divergent evolutionary trajectories, or shared due to their convergent adaptation to a common intracellular niche^3–5^. Nonetheless, our understanding of these species is still greatly limited by the lack of axenic culture systems and robust genetic tools that empower pathogen-directed investigations^6^.

These principles of obligate intracellular infection are well-exemplified by pathogens within *Rickettsia*, a Gram-negative genus responsible for a range of emerging and re-emerging arthropod-borne vascular diseases^7^ (*e.g.*, Rocky Mountain spotted fever). Within *Rickettsia,* the tick-borne spotted fever group (SFG) is increasingly recognized as a particularly useful model for how bacteria adapt to and infect the host cytosolic niche^2^. SFG rickettsial pathogens undergo an intracellular lifecycle that begins with host cell invasion followed by escape into the cytosol (**Fig. 1A**). There, SFG rickettsiae replicate and assemble tails of host actin that enable pathogen cytosolic motility, ultimately driving cell-to-cell spread and dissemination. Although the rickettsial lifecycle superficially resembles that of other intracellular pathogens like *Listeria monocytogenes*, SFG *Rickettsia* spp. exhibit divergent underlying mechanisms of host interaction, likely due to their highly minimized genomes and specialized evolutionary trajectory^2^. This includes unique interactions with host receptors^8^ and innate defense pathways^9^, as well as host cytoskeletal components that support actin-based motility and intercellular spread^10,11^. Despite recent progress in rickettsial genetic tools^12,13^, the traditional constraints of studying obligate intracellular pathogens are a limitation on our ability to probe how SFG *Rickettsia* spp. interact with and depend on their host cells. The lack of vaccines and effective diagnostics for SFG rickettsial infections underscores the need for new insights into the infection dependencies of these pathogens.

**Figure 1.**
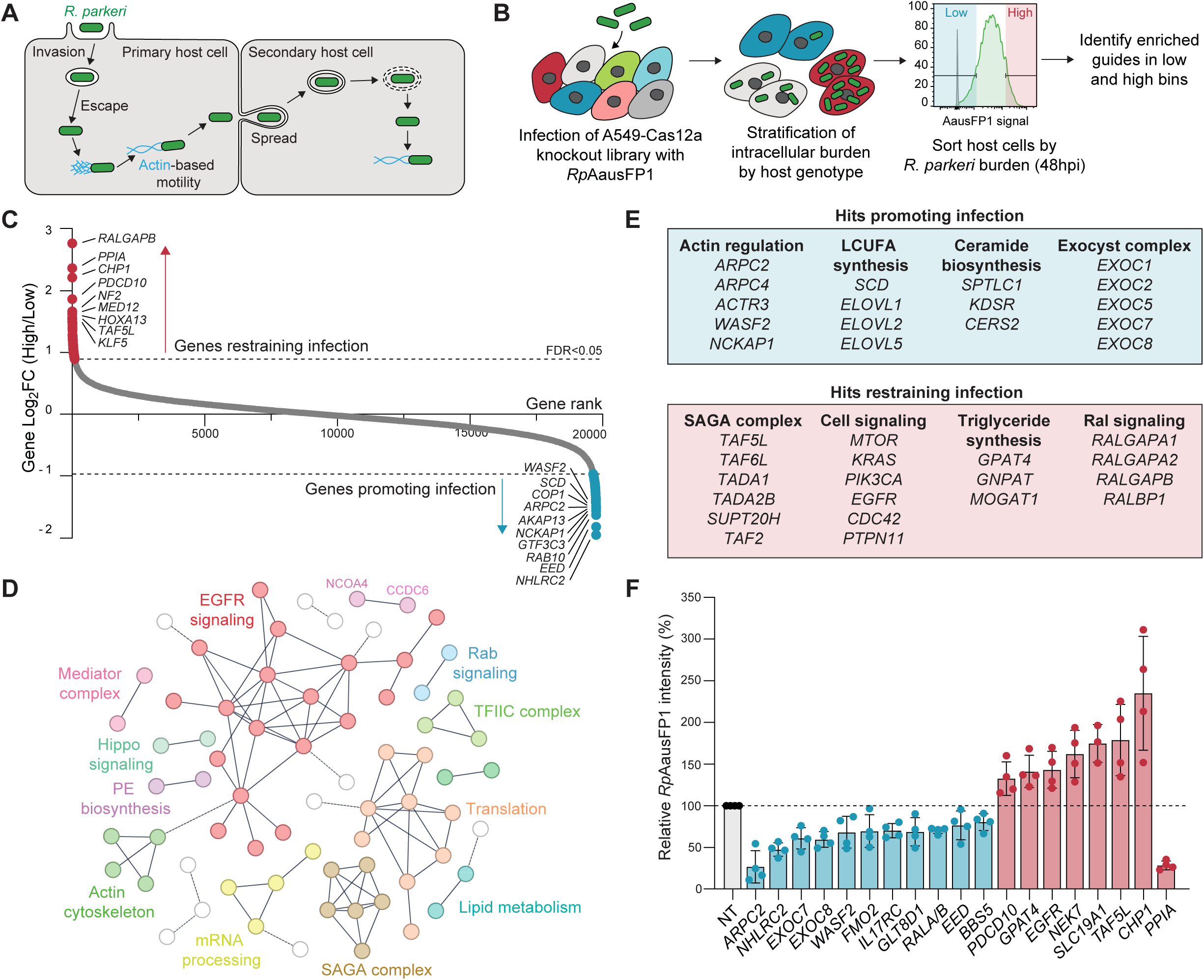
Genome-scale CRISPR/Cas12a screen for host determinants of *R. parkeri* infection. (**A**): Intracellular lifecycle of *R. parkeri* in mammalian cells. (**B**): Screen design and workflow. The top and bottom 10% fluorescence bins were sorted from infected libraries to identify human genes impacting *Rp*AausFP1 burden. (**C**): Waterfall plot of screen results. Mean log2 fold change (LFC) for gRNAs targeting each gene between screen replicates is plotted. Dashed lines correspond to LFC values with false discovery rates (FDR) <0.05. (**D**): Network analysis of hits with FDR <0.05. Hits without a STRING interaction confidence value > 0.9 to another hit (solid lines) were not visualized. Colors correspond to DBSCAN-identified clusters, and dotted lines additionally denote cluster edges. The NCOA4/CCDC6 cluster did not have a clear functional annotation. (**E**): Selected functionally related hits from cluster analysis and manual curation of screen results. All listed hits had screen LFC > 0.5 in the indicated direction. LCUFA: long-chain unsaturated fatty acids. (**F**): Hit validation with targeted enAsCas12a gene knockouts in A549s at 48 hpi with *Rp*AausFP1. Note *RALA/B* is a double knockout construct. Each point represents the relative geometric mean GFP intensity of a biological replicate compared to a matched non-target (NT) well processed on the same day. Error bars depict SD.

Host-directed approaches can query pathogen biology by perturbing its host environment, circumventing the need for extensive pathogen genetic manipulation and making them especially well-suited for studies of obligate intracellular bacterial infection. Such approaches, in particular genome-wide functional genetic screens, have been fruitful in dissecting microbial pathogenesis in other systems, for example host receptor tropism across a wide range of viruses^14^. However, these tools have only been sparingly deployed in targeted settings to study obligate intracellular bacteria^15–17^, potentially due to technical barriers to infecting large host cell libraries and inadequate screening-compatible functional readouts. Here, we developed a host-directed functional genomic screening platform to investigate obligate intracellular bacterial infection at scale, focusing on the representative SFG species *Rickettsia parkeri* as a model pathogen. This screen uncovered numerous novel host regulators of intracellular *R. parkeri* fitness comprising diverse functions and pathways. We found that a top screen hit, the host proline isomerase PPIA (cyclophilin A), is a critical and previously unrecognized host determinant of infection, and mechanistic dissection of this hit uncovered a unique surface protein maturation event tied to pathogenesis. We further show that this host dependency is not unique to *Rickettsia* but a convergent feature of the obligate intracellular lifestyle, deepening our understanding of host cell interactions with this understudied class of pathogen.

## Results

### Genome-scale identification of host determinants of *R. parkeri* fitness

To identify human genes that influence pathogen fitness across the intracellular *R. parkeri* lifecycle (**Fig. 1A**), we designed a flow cytometry-based, host-directed functional genetic screen. We first identified a suitable challenge strain by screening *R. parkeri* strains expressing various fluorescent proteins and found that *R. parkeri* expressing either TagBFP or the bright GFP variant AausFP1^18^ (*Rp*AausFP1) had superior fluorescence compared to established GFPuv-expressing *R. parkeri* strains (**Fig. S1A**). However, only *Rp*AausFP1 withstood methanol fixation, which can be preferrable to crosslinking fixation methods for sequencing-based methods^19^ (**Fig. S1B**). *Rp*AausFP1 was indistinguishable from WT *R. parkeri* in plaque assays and the fluorescence of infected host cells closely matched the signal from staining with an α-*Rickettsia* antibody (**Fig. S1C-E**). These data demonstrate that AausFP1 provides a faithful readout of *R. parkeri* burden and led us to select *Rp*AausFP1 for the screen.

We next used *Rp*AausFP1 to infect a genome-scale knockout library of A549 human epithelial cells stably expressing enAsCas12a (**Fig. 1B**). We reasoned that gRNAs enriched in the high or low fluorescence bins would represent host genes that normally restrain or promote *R. parkeri* fitness, respectively. We selected a Cas12a-based platform with tandem guide constructs as it lowers screen cell coverage requirements by ∼50% compared to Cas9^20^. At 48 hours post-infection (hpi), a timepoint where >95% of host cells were infected (**Fig. S1F-G** and **S2A**), the library was fixed and the top and bottom 10% of infected cells were isolated by FACS and sequenced to identify host loci impacting *R. parkeri* burden. Known essential genes were strongly depleted in the input library, confirming robust Cas12a-based editing in our system (**Fig. S2B**). In sorted samples, we also noted a low overall correlation between gRNA fold changes in the high versus low fluorescence bins, which is often observed in flow cytometry-based screens due to high selective pressure and cell number loss^21^ (**Fig. S2C**). By comparing gRNA fold changes in the high versus the low bins across two screen replicates, we identified 96 high and 113 low bin hits with corrected FDR < 0.05 (**Fig. 1C** and **Table S1**). Cluster and manual analyses highlighted the wide range of host processes required for optimal *R. parkeri* infection (**Fig. 1D-E** and **Table S2**). These included core host cell processes (*e.g*., the SAGA chromatin remodeling complex), metabolic pathways (*e.g.,* lipid biosynthesis) and intracellular trafficking mechanisms (*e.g.*, exocytosis). Importantly, three components of the Arp2/3 actin nucleation complex (*ARPC2, ARPC4,* and *ACTR3*), were identified as promoting infection. Arp2/3 is known to facilitate *R. parkeri* invasion^22^, validating the overall screening approach. We noted opposing phenotypes for different lipid metabolic pathways (long-chain unsaturated fatty acid, ceramide, and triglyceride biosynthesis) (**Fig. 1E**), suggesting that competition between distinct arms of host lipid metabolism is a key regulator of intracellular *R. parkeri* fitness. This concept is consistent with prior targeted work in other cell types^23^ and highlights the ability of the screen to identify functionally concordant hits.

For hit validation, we generated single gene knockout A549 lines with enAsCas12a and infected them with *Rp*AausFP1. To enrich for host loci that directly regulate *R. parkeri* as opposed to exerting general effects on host cell fitness that could indirectly impact pathogen burden, we focused on hits that were not already strongly selected in the input (**Fig. S2D**). As expected, knockout lines for hits in the low infection bin (*i.e.*, required for infection) exhibited lower levels of *Rp*AausFP1 than non-target (NT) controls, and *vice versa* (**Fig. 1F**). Some validated hits have not previously been described to contribute to rickettsial infection but have known roles in intracellular bacterial pathogenesis (*e.g.*, *EXOC7/8*^24^ and *NHLRC2*^25^). However, we also validated hits that have not been associated with any other mammalian bacterial or microbial infection (*e.g., NF2* and *CHP1*). Surprisingly, one of the top hits from the screen, *PPIA*, exhibited an opposing phenotype to the direction of its original bin (**Fig. 1F**, rightmost). This unexpected observation prioritized *PPIA* for further investigation.

### *PPIA* is essential for *R. parkeri* intercellular spread and actin-based motility

PPIA, also known as cyclophilin A or CypA, is a widely conserved *cis-trans* peptidyl-prolyl isomerase with numerous reported roles in host cell signaling and biology^26^. PPIA has been implicated in host susceptibility to diverse viral infections, particularly HIV-1, but its role in bacterial pathogenesis is not well understood^27^. No other members of the cyclophilin family were identified as hits and single knockouts of every other major human cyclophilin (*PPIB/C/D/E/F/G/H*) had no effect on infection, indicating PPIA has a specific contribution to *R. parkeri* fitness (**Fig. S3A**). In flow cytometry of infected *PPIA*ko cells, we noted an approximately bimodal phenotype with a distinct subpopulation of hyper-infected cells, contrasting with the uniformly right-shifted phenotype of other “high” hits (*e.g.*, *CHP1*) (**Fig. 2A-B**). We hypothesized that this phenotype could be explained in the screen by reduced pathogen spread, leading to buildup of *R. parkeri* in a small number of *PPIA*ko cells that would move into the high fluorescence bin relative to cells with unrelated edits. Consistent with this idea, we observed a near-complete loss of pathogen spread in fluorescence microscopy of host cell monolayers sparsely infected with *R. parkeri* (**Fig. 2C-D**). Importantly, the bacterial number per cluster of infected host cells was unaltered, indicating PPIA is not required for *R. parkeri* replication (**Fig. 2E**). Re-expression of WT PPIA, but not a mutant lacking the key catalytic and substrate binding pocket residue^28^ Arg55 (PPIA^R55A^), completely rescued the spread phenotype (**Fig. 2C-D**), indicating that PPIA’s effect on *R. parkeri* infection requires its capacity to bind and isomerize substrates.

**Figure 2.**
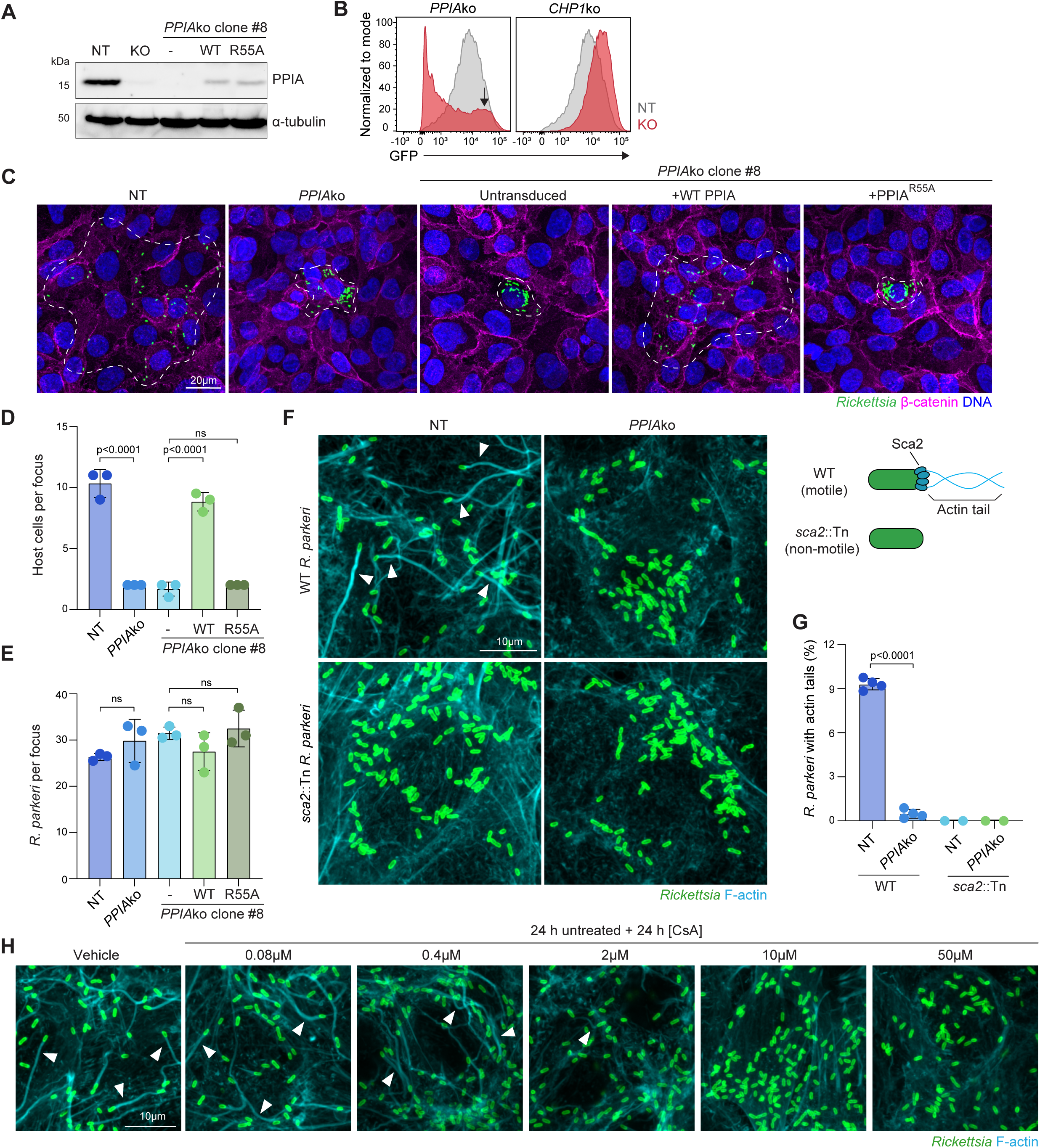
Host PPIA is required for *R. parkeri* spread and actin-based motility. (**A**): Western blot confirming loss of PPIA in pools (KO) and clonal cell lines with rescue constructs. Data are representative of two independent blots with cells from different passages. (**B**): Flow cytometry curves of *CHP1*ko and *PPIA*ko cells infected with *Rp*AausFP1 at 48 hpi. Grey curves represent matched NT samples (same between knockouts). Curves are representative of n=3 biological replicates. Arrow indicates hyper-infected population in *PPIA*ko cells. (**C**): Spread assay of WT *R. parkeri* in NT and *PPIA*ko cell lines. Dashed lines outline representative infection foci. (**D** and **E**): Quantification of spread assays for number of host cells (D) and bacteria per focus (E). Means and SD from n=2-4 biological replicates are plotted. Means were used to calculate p-values (one-way ANOVA with Holm-Šídák multiple comparisons test). (**F**): Actin tails at 48 hpi in NT and *PPIA*ko cells infected with WT or *sca2*::Tn *R. parkeri*. Expected strain phenotypes are diagrammed on the right for reference. (**G**): Quantification of data from Panel F. Means and SD from n=3 biological replicates (>2000 bacteria counted per replicate) are plotted. Means were used to calculate p-values (Welch’s t-test). (**H**): Actin tails at 48 hpi in cells treated with varying concentrations of cyclosporin A (CsA) for 24 h, beginning at 24 hpi. Arrowheads highlight bacteria with actin tails. Images are representative of at least 3 biological replicates.

The lack of *R. parkeri* spread in *PPIA*ko host cells resembles the known phenotype of *R. parkeri* lacking Sca2 (*sca2*::Tn), the pathogen surface protein primarily responsible for actin tail nucleation and cytosolic motility^29,30^. Indeed, actin tails were completely absent in *R. parkeri-*infected *PPIA*ko host cells, phenocopying the loss of actin tails seen with *sca2*::Tn infections (**Fig. 2F-G**). Actin tail formation was also blocked when infected cells were treated with the FDA-approved PPIA inhibitor cyclosporin A (CsA) or NIM811, a non-immunosuppressive CsA analog^31^, orthogonally supporting a role for PPIA in this stage of infection (**Fig. 2H** and **S3B**). Inhibitors of the other two recognized proline isomerase families, FK506 (FKBPs) and juglone (parvulins), did not affect actin tails (**Fig. S3B**), indicating that proline isomerization in general is not required for *R. parkeri* actin-based motility. From these data, we concluded that host PPIA is essential for *R. parkeri* actin tail formation and subsequent cell-to-cell spread.

### PPIA localizes at the *R. parkeri* actin-associated pole

To further understand the role of PPIA in actin-based motility, we asked where PPIA was localized during infection. We adopted a previously established approach from studies of PPIA localization to the HIV-1 capsid using PPIA fused to the tetrameric RFP derivative dsRed^32,33^. We modified this construct by replacing dsRed with the less cytotoxic variant dsRed-Express2 (dRE2)^34^. Strikingly, during *R. parkeri* infection of WT A549 cells harboring the fusion construct, we observed polar localization of PPIA-dRE2 in approximately 12% of bacteria at 48 hpi (**Fig. 3A-B**). Polar PPIA-dRE2 signal was associated with *R. parkeri* with and without actin tails, indicating PPIA does not require actin to localize to the bacterium. There was also no apparent PPIA-dRE2 enrichment along the actin tail itself or host cell stress fibers. Interestingly, in the subset of *R. parkeri* that had both polar PPIA-dRE2 signal and an actin tail, dRE2 signal localized to the actin-associated pole of the bacterium 97% of the time (88 out of 91 scored bacteria). Polar PPIA-dRE2 localization was lost in infected cells expressing PPIA^R55A^-dRE2 (**Fig. 3A-B**) or treated with CsA (**Fig. 3C-D**), suggesting that localization is dependent on PPIA activity and/or substrate binding. PPIA’s polar localization raised the possibility that it targets a substrate at the bacterial pole that contributes to actin-based motility. Since the *R. parkeri* actin-nucleating factor Sca2 is enriched at the actin-associated pole^29^, we hypothesized Sca2 may be the target of PPIA. Accordingly, PPIA-dRE2 failed to localize to *R. parkeri* when cells were infected with *sca2*::Tn bacteria (**Fig. 3E-F**), implicating Sca2 as the mediator of PPIA recruitment to the *R. parkeri* pole.

**Figure 3.**
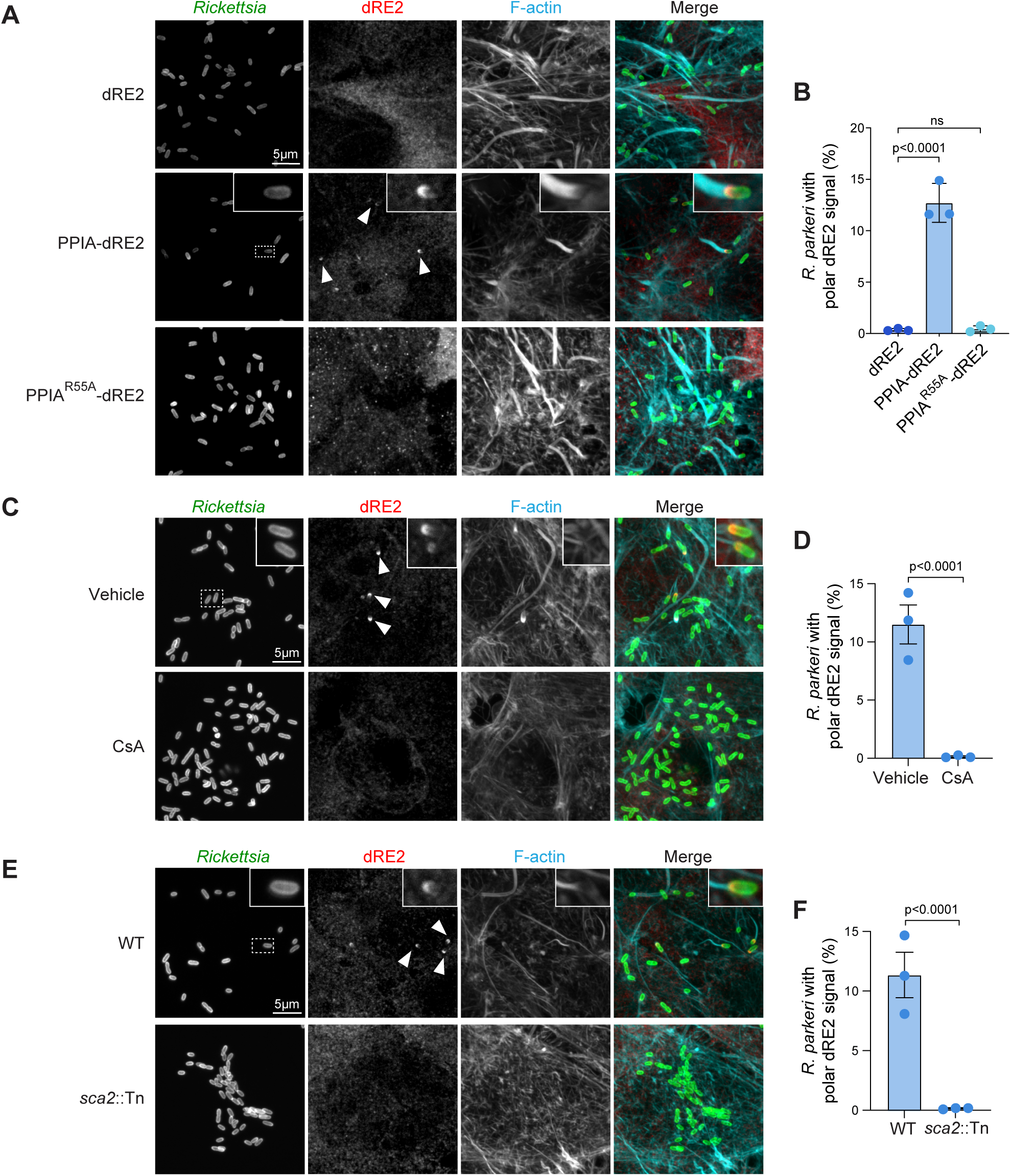
PPIA localizes to the *R. parkeri* actin-associated pole in a Sca2-dependent manner. (**A**): Localization of dsRed-Express2 (dRE2) constructs in clonal stably transduced A549s infected with WT *R. parkeri* at 48 hpi. Arrowheads highlight bacteria with polar PPIA-dRE2 signal. (**B**): Quantification of polar dRE2 signal frequencies from experiments in Panel A. (**C**): PPIA-dRE2 localization in infected A549s treated with 10μM CsA for 24 h at 24 hpi. (**D**): Quantification of data in Panel C. (**E**): PPIA-dRE2 localization in infections with *sca2*::Tn *R. parkeri.* (**F**): Quantification of data in Panel E. In B, D and F, means and SD from n=3 biological replicates are plotted (>1000 bacteria counted per replicate). Means were used to calculate p-values (one-way ANOVA with Holm-Šídák multiple comparisons test (B) or a Welch’s t-test (D and F)). Images are representative of n=3 biological replicates.

### PPIA interacts directly with the Sca2 passenger domain C-terminus

Noting the co-occurrence of Sca2 and PPIA at the actin-associated pole, we next hypothesized that the two proteins directly interact. Sca2 belongs to the autotransporter family of bacterial outer membrane proteins, which consist of an N-terminal passenger domain that is extruded to the surface by transit through a C-terminal β-barrel^35^ (**Fig. 4A**). The Sca2 passenger domain has an atypical globular α-helical structure, rather than the β-solenoids that characterize model autotransporters (**Fig. S4A**). The domain comprises N-terminal sequences that mediate actin nucleation and a C-terminal region containing a C-repeat domain (CRD) with no clearly defined function^36–39^. AlphaFold3 modeling of a PPIA-Sca2 interaction using the full *R. parkeri* Sca2 sequence revealed a high-confidence (iPTM=0.73) predicted interaction localized to the Sca2 C-terminal region near the CRD (**Fig. 4B-C**). To test this prediction, we purified GST-tagged truncation constructs of *R. parkeri* Sca2 and used them for *in vitro* pulldowns with purified PPIA (**Fig. 4D**). Sca2(34-1544, PD) containing the entire passenger domain pulled down PPIA. Sca2(34-1086, PDΔ), which lacks the C-terminal region, did not pull down PPIA, whereas a construct containing only this stretch of sequence did (Sca2(1086-1544), CRD) (**Fig. 4D**). These results demonstrate the necessity and sufficiency of the Sca2 CRD-containing region for PPIA binding.

**Figure 4.**
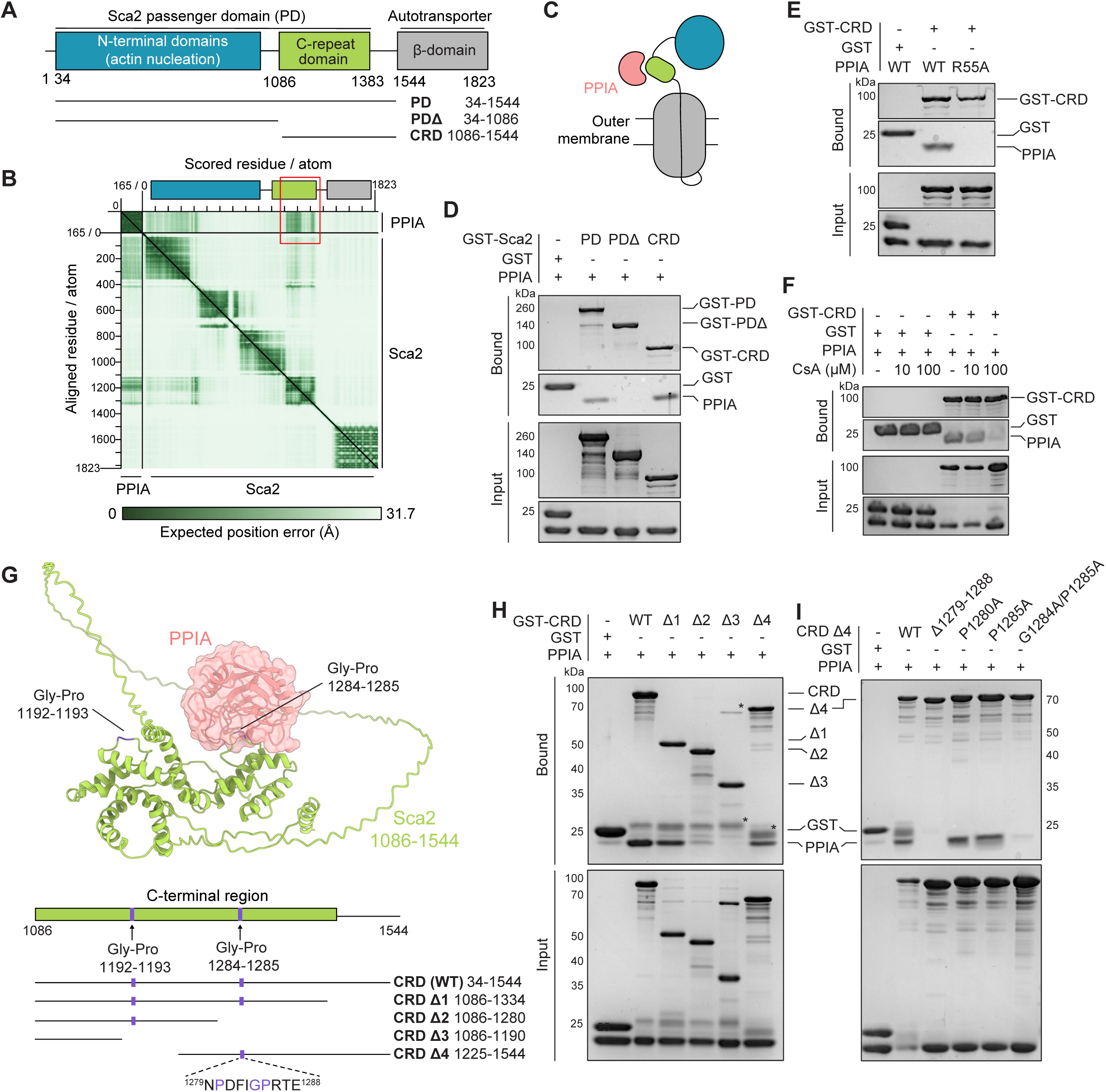
PPIA interacts directly with the C-terminal passenger region of *R. parkeri* Sca2. (**A**): Domain diagram of *R. parkeri* Sca2. (**B**): Position-adjusted error (PAE) plot of the AlphaFold3 model of the full-length Sca2 and PPIA interaction. The region where the two proteins are predicted to interact is boxed in red. (**C**): Simplified diagram of Sca2 topology and predicted PPIA binding location. (**D-F**): Glutathione pulldowns of PPIA with Sca2 domain constructs indicated in panel A (D), WT Sca2 and PPIA^R55A^ (E), or with 10-100 μM cyclosporin A (CsA) in the binding reaction (F). (**G**): Top: predicted PPIA-Sca2 (1086-1544) interaction. The two Gly-Pro sites are indicated in orange and PPIA is surface shaded. Bottom: schematic of C-terminal region and locations of the two Gly-Pro sites. (**H-I**): Glutathione pulldown with WT PPIA and Sca2 CRD truncations (H) or specific mutants of CRD Δ4 indicated in panel G (I). * = non-specific band or possible spontaneous GST cleavage product. All images shown are Coomassie blue stained SDS-PAGE gels representative of at least 2 independent pulldowns.

Canonical PPIA binding occurs via its main catalytic pocket, which contains several well-characterized residues that mediate both substrate binding and isomerase activity^40^. Of these, Arg55 is the most often targeted in loss-of-function studies, and we earlier demonstrated that this residue is required for PPIA’s control of *R. parkeri* spread (**Fig. 2C-D**). We found that purified PPIA^R55A^ could not be pulled down by Sca2(1086-1544) (**Fig. 4E**) and addition of the competitive PPIA inhibitor CsA, which binds the catalytic pocket, also blocked pulldown of WT PPIA (**Fig. 4F**). We concluded from these data that PPIA binds Sca2 through its primary binding pocket, suggesting that Sca2 is an isomerization substrate of PPIA.

PPIA binds a wide range of substrates with reported preference for sequences containing Gly-Pro dipeptides^41^. Sca2(1086-1544) harbors two Gly-Pro sequences in the CRD (residues 1192-1193 or 1284-1285), which are largely conserved in SFG *Rickettsia* spp. as well as species in other *Rickettsia* subgroups (**Fig. S4B**). Moreover, none of the known proline-rich sequences in the Sca2 N-terminal region^36^ were sufficient for the interaction (**Fig. 4D**). Using AlphaFold3 to model the interaction between PPIA and the CRD, we observed that PPIA was predicted to interact with the second Gly-Pro position at amino acids 1284-1285 (iPTM=0.85) (**Fig. 4G, S4C**). This dipeptide lies in a flexible loop comprising Sca2(1279-1288) that extrudes into the PPIA catalytic pocket, where it makes contacts with critical PPIA residues including Arg55^28^ (**Fig. S4D-E**). This model closely resembles the experimentally determined structure of PPIA’s interaction with the HIV-1 capsid protein^42,43^ (**Fig. S4E**). Consistent with this prediction, an additional series of CRD truncations demonstrated that Gly1284-Pro1285, and not Gly1192-Pro1193, is required for optimal PPIA binding (**Fig. 4H**). Although deletion of the entire loop abolished binding, single mutations to the two prolines within this loop, Pro1280 or Pro1285, had no effect on binding (**Fig. 4I**). We only observed loss of binding when we mutated both Gly1284-Pro1285 (**Fig. 4I**), suggesting that the PPIA-Sca2 CRD interaction may involve distinct sequence requirements in the loop beyond Pro1285.

### Sca2 surface exposure requires host cell PPIA

In model autotransporters, the passenger domain C-terminus is thought to exit the β-barrel as a hairpin and undergo an early extracellular folding event critical for translocation of the remainder of the passenger domain^44–48^. Our finding that PPIA directly binds the Sca2 passenger C-terminal region therefore suggests that PPIA acts at the level of Sca2 surface exposure, explaining its requirement for actin tail formation. To test this idea, we leveraged our standard immunofluorescence staining protocol, which does not permeabilize *R. parkeri* and is thus surface-selective^49^. Using a validated α-Sca2 antibody raised against the first 500aa of the passenger domain^36^, we found that surface Sca2 was lost in *R. parkeri*-infected *PPIA*ko cells at 48 hpi (**Fig. 5A**, left and **Fig. 5B**). Bacteria lacking surface Sca2 still produced Sca2 protein, as subsequent permeabilization of *R. parkeri* with lysozyme revealed comparable total frequency and levels of Sca2 between *PPIA*ko and NT infections (**Fig. 5A**, right, and **Fig. 5B-C**). Total Sca2 levels were also unaltered in immunoblots of infected cells (**Fig. S5A**), suggesting PPIA does not regulate the production of Sca2 within *R. parkeri*. Treatment of WT A549 cells with CsA or NIM811 blocked Sca2 surface exposure (**Fig. 5D**), independently confirming the role of PPIA. To assess the dynamic nature of this process, we performed a washout experiment where we removed CsA at different times during infection. As little as 2 hr of CsA washout enabled recovery of Sca2 surface exposure and actin tails (**Fig. S5B-C**), suggesting that Sca2 translocation can resume or is continually produced during infection. Together, these findings support a model where PPIA binds the Sca2 C-terminal passenger region during surface translocation, promoting its stabilization or folding and thus enabling passenger domain surface exposure (**Fig. 5E**).

**Figure 5.**
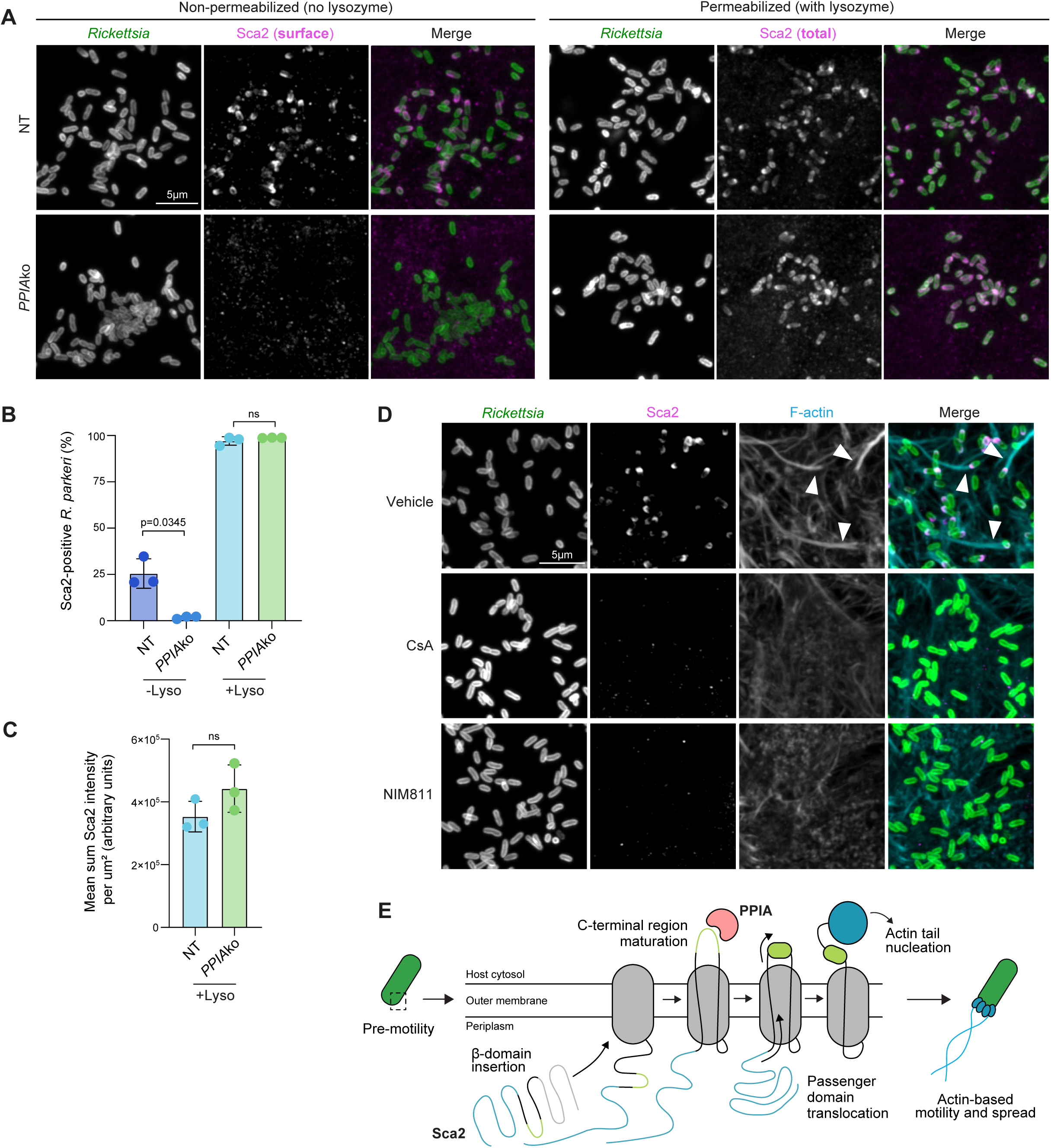
PPIA is required for surface exposure of Sca2. (**A**): Sca2 staining in lysozyme-treated or untreated NT or *PPIA*ko A549 cells infected with WT *R. parkeri* at 48 hpi. Left: non-permeabilized (surface staining only) samples. Right: permeabilized (total staining) samples. (**B**): Quantification of Sca2-positive bacteria in lysozyme-treated or untreated samples. (**C**): Quantification of Sca2 intensity in lysozyme-treated samples. Samples were normalized by dividing by the area of each bacterium.(**D**): Effect of 10μM CsA or NIM811 treatment on Sca2 surface levels during infection of WT A549 cells at 48 hpi. Arrowheads highlight bacteria with actin tails. Means and SD from n=3 biological replicates are plotted (>2000 bacteria counted per replicate). Means were used to calculate p-values (Welch’s t-test). Images are representative of n=3 biological replicates. (**E**): Proposed model of PPIA-enabled Sca2 surface translocation. PPIA binds the Sca2 CRD as it is exposed to the extracellular space, stabilizing, or folding it and enabling extrusion of the remainder of the passenger domain, ultimately allowing actin tail nucleation. In the absence of PPIA, this process likely stalls at hairpin formation and the passenger domain is never exposed.

### PPIA shapes *R. parkeri* infection across cellular and *in vivo* contexts

Sca2-dependent actin-based motility has been observed across a variety of eukaryotic cell types, prompting us to ask whether dependence of this process on PPIA was conserved across the different cellular contexts experienced by *R. parkeri*. Actin tails were completely absent in *PPIA*ko human vascular endothelial-like cells (EA.hy926) (**Fig. 6A-B**), demonstrating this phenotype is conserved in the physiological target cell type of *Rickettsia* spp.^50,51^. Similarly, when we treated a model *Ixodes scapularis* tick cell line (ISE6) with NIM811, we observed a complete loss of actin tails (**Fig. 6C**), extending the relevance of PPIA to the arthropod vector environment. We orthogonally validated these results with CsA treatment in both cell lines (**Fig. S6A-B**), underscoring that PPIA is likely a universal requirement for *R. parkeri* dissemination across cell types.

**Figure 6.**
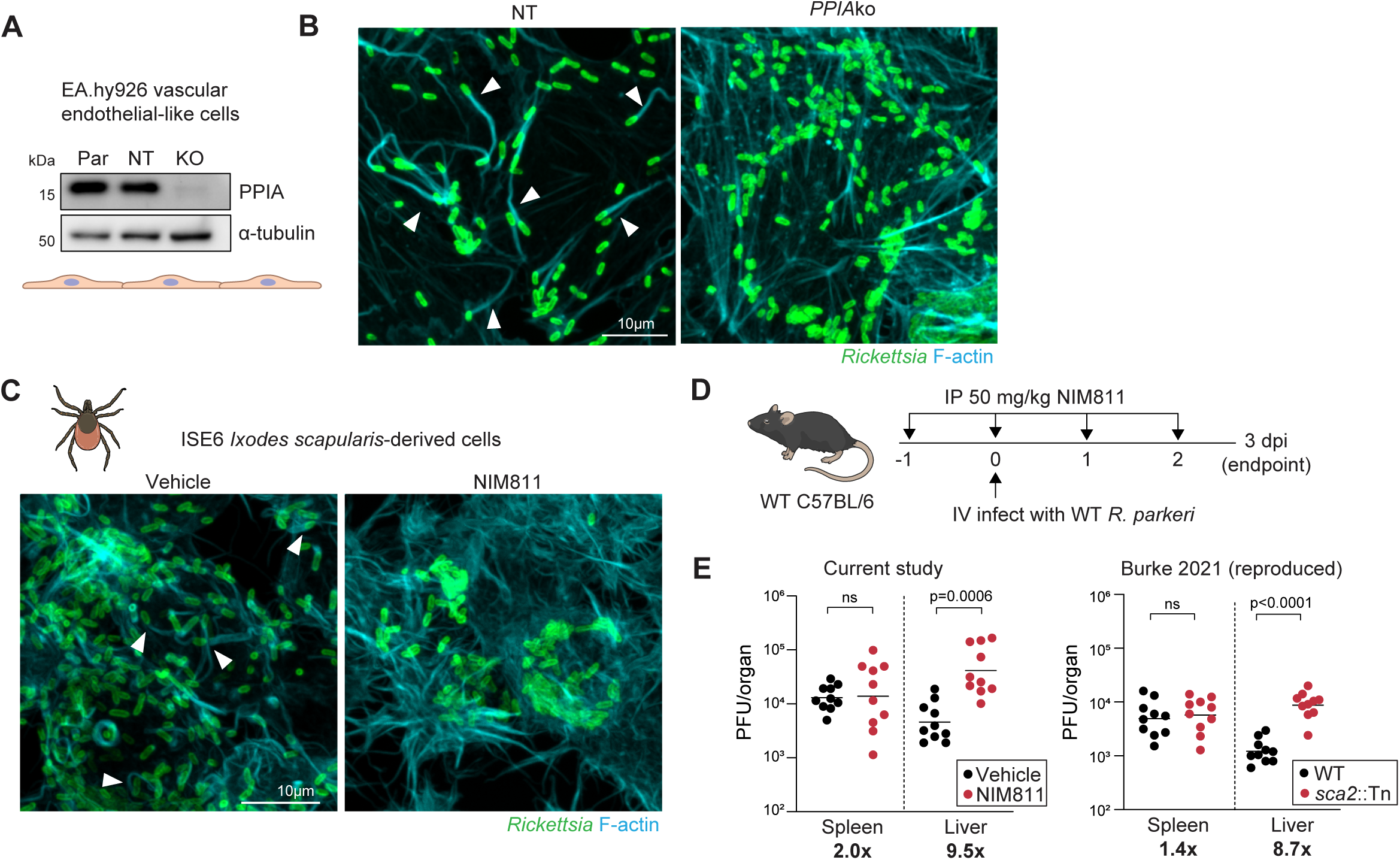
PPIA shapes *R. parkeri* infection across cellular and *in vivo* contexts. (**A**): Western blot of parental (Par), non-target (NT) and *PPIA*ko (KO) EA.hy926 vascular endothelial cells. (**B**): Actin tails at 48 hpi in NT or *PPIA*ko EA.hy926 cells. (**C**): Actin tails at 48 hpi in *Ixodes scapularis* ISE6 cells treated with vehicle (DMSO) or 5 μM NIM811 starting at 1 hpi. (**D**): Schematic of mouse infection experiment. (**E**): *R. parkeri* plaque-forming units (PFU) in spleen (left) and livers (right) from infected mice. The right graph shows the known liver-specific phenotype of *sca2*::Tn *R. parkeri* at 3 dpi in i.v.-infected WT C57BL/6 mice (data reproduced from (Burke *et al*, *eLife* 2021)). Each point represents one mouse. Data were aggregated from two independent experiments and analyzed with Mann-Whitney U tests. All images are representative of n=3 biological replicates infected with WT *R. parkeri*. Arrowheads highlight bacteria with actin tails.

Sca2 is a *bona fide* SFG rickettsial virulence factor, and *sca2*-deficient bacteria fail to elicit robust disease in multiple animal models^52,53^. To examine whether inhibition of PPIA would phenocopy the loss of Sca2 in an *in vivo* setting, we used an intravenous *R. parkeri* infection model in WT C57BL/6 mice^53^ (**Fig. 6D**). In this model, *sca2*::Tn *R. parkeri* exhibit elevated bacterial burdens in the liver, potentially because they cannot disseminate effectively^53^. We asked whether a host-directed PPIA inhibitor could recapitulate this phenotype. For *in vivo* studies, we chose NIM811 because CsA has well-known immunosuppressive effects unrelated to PPIA enzymatic activity^54,55^. Intraperitoneal administration of NIM811 in infected mice closely phenocopied the *sca2*::Tn liver-specific increase in burdens (**Fig. 6E**). Importantly, NIM811 did not significantly increase liver burdens in mice infected with *sca2*::Tn *R. parkeri* (**Fig. S6C-D**), demonstrating that the effect is Sca2-dependent and that the PPIA-Sca2 axis facilitates infection *in vivo*.

### Shared dependency on host cyclophilins in obligate intracellular bacteria

Cyclophilin proline isomerases are conserved across all domains of life. While most Gram-negative bacteria, including *E. coli*, encode periplasmic and cytoplasmic cyclophilins^56^, evidence of cyclophilin secretion in the intracellular pathogen *Legionella pneumophila* suggests that external proline isomerase activity also contributes to bacterial fitness^57^. The surface dependence of Sca2 on PPIA suggests that species like *R. parkeri* may lack endogenous cyclophilin activity, instead relying on host-supplied cyclophilins to facilitate key virulence behaviors like surface protein maturation. Accordingly, we found that the cyclophilin family (cyclophilin type peptidyl-prolyl cis-trans isomerase/CLD, PF00160) is completely absent from not only *R. parkeri*, but all *Rickettsia* spp., despite broad conservation across their class (Alphaproteobacteria, 89.8% (8681 of 9662) genomes) (**Fig. 7A, Table S4**). Strikingly, cyclophilins were also absent in many other known obligate intracellular bacterial pathogens, including those not dependent on actin-based motility for infection (*e.g., Orientia* and *Chlamydia*). In contrast, PF00160 was only absent from one genus of facultative intracellular pathogens (*Francisella*). This conservation pattern was restricted to cyclophilin-like domains and not proline isomerases in general, as predicted FKBP-type (PF00254) and parvulin-type (PF13145) isomerases are present (**Fig. 7A**) and have been biochemically characterized in *Rickettsia* spp.^58^. Therefore, the specific absence of cyclophilin-type proline isomerases may be associated with obligate intracellular bacterial lifestyles.

**Figure 7.**
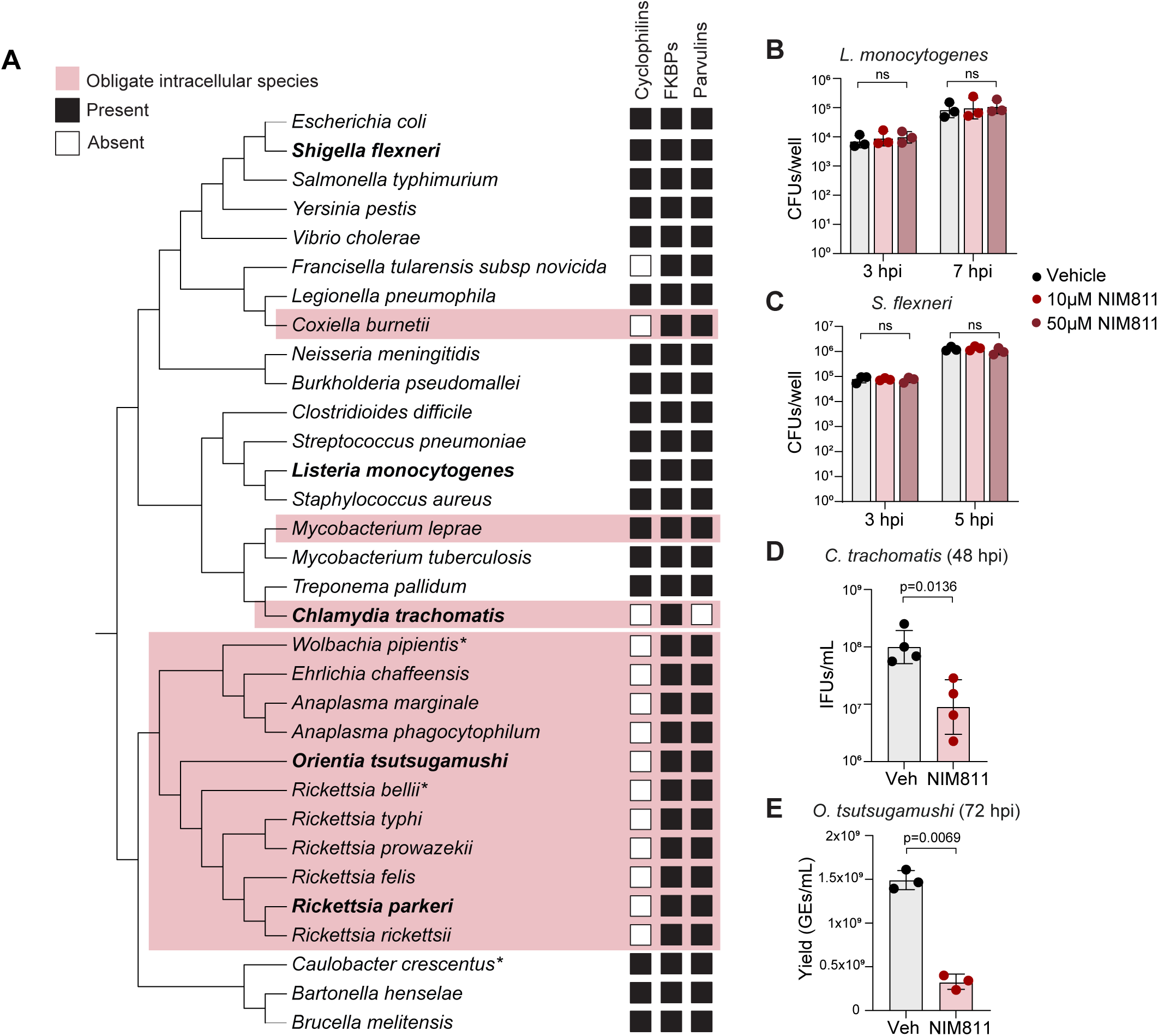
Shared dependencies on host cyclophilins in obligate intracellular bacterial pathogens. (**A**): Cladogram of selected bacterial species representative of extracellular and facultative or obligate intracellular lifestyles. Species not known to be pathogenic to humans are marked with *. Species shaded in pink are obligate intracellular microbes. Branch lengths do not represent distances. Open and filled squares indicate the absence or presence, respectively, of each proline isomerase family. Strains included in the analysis are listed in Table S3. (**B** and **C**): Colony-forming units (CFUs) of *L. monocytogenes* (B) and *S. flexneri* (C) at the indicated timepoints in A549 cells treated with vehicle or the indicated concentration of NIM811 at 1 hpi. (**D**): Inclusion-forming units (IFUs) of *C. trachomatis* at 48 hpi in HeLa cells treated with vehicle (Veh) or 10 μM NIM811 at 5 hpi. (**E**): Input-adjusted gequivalents (GEs) of *O. tsutsugamushi* at 72 hpi in EA.hy926 cells treated with vehicle or 10 μM NIM811 at 4 hpi. Means and SD from at n=3-4 biological replicates are plotted (geometric means in D and E). Means were used to calculate p-values (one-way ANOVA with Holm-Šídák multiple comparisons test (B and C) or lognormal Welch’s t-test (D and E)).

To experimentally test whether the absence of a PF00160 homolog correlates with an infection dependency on host cyclophilins, we evaluated whether NIM811 affected the intracellular fitness of two facultative and two obligate intracellular bacterial pathogens. Addition of NIM811 after pathogen entry had no discernable effect, even at very high doses, on the intracellular replication of *L. monocytogenes* or *S. flexneri,* which both encode endogenous cyclophilins (**Fig. 7B-C**). In contrast, NIM811 significantly reduced the expansion of two distinct obligate intracellular pathogens *C. trachomatis* and *O. tsutsugamushi* (**Fig. 7D-E**). This bifurcation of NIM811 sensitivity indicates that host cyclophilin dependency is a previously unrecognized common feature of obligate intracellular bacterial infection.

## Discussion

The unique host-bacterial interactions arising from the extreme biological constraints imposed by the obligate intracellular lifestyle have remained largely unexplored. Through a host-directed functional genetic screen, we revealed the first large-scale dataset of host determinants of SFG rickettsial infection, subsequently identifying direct surface control of the *R. parkeri* actin-nucleating virulence factor Sca2 by the host proline isomerase PPIA. Host-directed maturation of a bacterial surface protein is a previously unrecognized mode of host-pathogen interaction. Dependency on PPIA is not specific to *R. parkeri*, as other obligate, but not facultative, intracellular bacteria required PPIA for robust infection. Thus, PPIA represents a broadly conserved host dependency of obligate intracellular bacteria whose varied contributions to diverse infectious life cycles will reveal new facets of how obligate pathogens have co-opted the eukaryotic cell.

The frequent loss of endogenous cyclophilins in obligate intracellular bacteria suggests that microbes that are highly adapted to the host cytoplasm have converged on PPIA as a dependency for productive infection. We speculate this shared requirement may have arisen due to PPIA’s exceptionally high intracellular abundance (∼16 µM in HEK293 cells^59^) and broad expression across tissues^60,61^. Importantly, PPIA dependencies are independent of the ability of a pathogen to form actin tails, as we found that multiple species that exhibit other forms of actin-based motility^62^ were not sensitive to a PPIA inhibitor. Sca2 is a *Rickettsia-*specific factor and neither *C. trachomatis* nor *O. tsutsugamushi* use actin-based motility, indicating that their PPIA dependencies will likely represent new molecular mechanisms of host-bacterial infection. Since *C. trachomatis* is vacuolar, in contrast to *R. parkeri* and *O. tsutsugamushi* which inhabit the cytosol, understanding convergent host dependencies like PPIA that may complement missing genomic functions could reveal additional, unexpected links between bacteria with otherwise divergent intracellular lifestyles and evolutionary trajectories.

PPIA bidirectionally influences infection by viruses including HIV-1, HCV, and influenza by binding to viral proteins and shaping their function(s)^27^. Our work therefore reinforces conceptual and biological parallels between obligate intracellular bacteria and viruses, which are both completely dependent on the host cell for survival. The viral dependency on host PPIA has long been considered an antiviral therapeutic target, and our findings warrant additional work to establish this concept in obligate intracellular bacterial infection. Such studies may be potentiated by the prior development of numerous cyclophilin inhibitors^31^ and clinical efforts in viral settings, including NIM811^63^. The concept of host-targeted therapies in SFG rickettsial infection has mainly been examined through the myriad predicted metabolic dependencies of these species^5,64^, for example host-provided metabolites like glutathione^3^ and isoprenoids^4^. It will be an interesting topic of future study to examine whether co-targeting dependencies with unique modes of action may elicit synergistic disruption of rickettsial infection.

The PPIA-Sca2 interaction is a conceptually distinct mode of post-translational virulence factor control that acts at the level of surface protein translocation, in contrast to typical regulatory mechanisms like altered protein stability or modifications like phosphorylation. Sca2-mediated actin tails are temporally regulated, but Sca2 levels do not similarly shift over the course of *R. parkeri* infection^29,65^, hinting its regulation has been partially outsourced on a post-translational level to a host element like PPIA. Determining whether key PPIA regulatory modifications like glutathionylation^66^ or acetylation^67^ can tune its activity and influence Sca2 maturation across cell types will be crucial for investigating host-dependent regulation of rickettsial actin-based motility. It remains an open question of whether PPIA may also participate in regulation of RickA, the complementary SFG rickettsial actin-based motility factor that transiently assembles tails in early infection by an orthogonal mechanism to Sca2^29,68^. Tissue-specific targeting of PPIA may also address the open question of how Sca2-dependent actin-based motility promotes *in vivo* SFG rickettsial dissemination and why the route of infection in mouse models has opposing effects on pathogen organ burdens^53^. Although PPIA can be secreted, circulating concentrations (∼13 nM in normal human plasma^69^) are ∼1000-fold lower than intracellular levels. This imbalance raises the possibility that PPIA could act as a host cytoplasmic signal for *R. parkeri* to activate and subsequently tune Sca2-dependent actin-based motility. Evaluation of this idea in other species could fundamentally widen the scope of intracellular bacterial cytoplasmic sensing mechanisms, which canonically rely on non-proteinaceous host-associated inputs like temperature^70^, pH^71^, and metabolites^72–74^.

Prior reports of PPIA in other bacterial infections have involved indirect host-mediated effects^75,76^ or direct interactions with secreted bacterial toxins^77–79^ or effector proteins^80–82^, all without PPIA recruitment to the pathogen itself. Our work identifies the first instance of a host cyclophilin acting directly on the bacterial surface, a new interaction site in an emerging model of PPIA as a central mediator of microbial infection outcome^27^. *Trans*-acting extracellular folding or isomerization events such as that likely exerted by PPIA on Sca2 have not been described for any other autotransporter, which are often bacterial virulence determinants. The prevailing model of autotransporter biogenesis involves the canonical β-solenoid passenger domain structure templating its own surface translocation^35^. While >90% of known autotransporters are β-solenoidal^83^, key autotransporters in pathogens like *Pseudomonas aeruginosa* (*e.g.*, EstA^84^) adopt α-helical passenger domain folds. The translocation mechanism of non-canonical passenger domains has yet to be explored. The Sca2 C-terminal domain likely contains an α-helical bundle^37^, but there is no experimental evidence to support the notion that it is self-templating. Our data favors a model where α-helical autotransporters like Sca2 follow a similar pathway to the surface as their β-solenoidal counterparts, but with additional dependencies on external factors, specifically cyclophilins. In free-living bacteria that do not have access to host cyclophilins, these mechanisms may be supported by secretion of endogenous cyclophilins, as has been reported in *L. pneumophilia*^85^. Sca2 is an excellent model to continue exploring this idea, as it is one of the few α-helical autotransporters with a known molecular function. The precise folding, stabilization and/or isomerization events in the PPIA-Sca2 interaction remain an open question, with implications for understanding the biogenesis and maturation of autotransporters with atypical passenger domains.

Lastly, our study demonstrates the potential of large-scale untargeted host-directed perturbations to access unexplored features of obligate intracellular bacterial biology. We anticipate that other hits from the *R. parkeri* screen comprise additional shared and unique infection dependencies. The mechanistic basis of the dozens of remaining screen hits will depend whether they act indirectly or directly on the pathogen, where in the cell they act, and whether they shape pathogen fitness at only one stage of the lifecycle or throughout infection. It remains to be seen whether a similar or divergent set of genes support *R. parkeri* infection across the cell types the pathogen encounters during infection (*e.g.,* vascular endothelial and innate immune cells^86^), a question that may be resolved by expanded genome-scale screening efforts to incorporate relevant host and pathogen contexts. Large-scale host-directed perturbations applied to currently intractable or poorly tractable bacterial pathogens will likely be instrumental to mine untapped bacterial diversity, revealing novel traits associated with bacterial adaptation to the intracellular space.

## Methods

### Eukaryotic cell lines

Human A549 lung epithelial cells, HEK293T human embryonic kidney, and African green monkey Vero kidney epithelial cell lines were obtained from the UC Berkeley Cell Culture Facility (Berkeley, CA, USA). Human EA.hy926 vascular endothelial cells were a gift from Daniel Swale (University of Florida, USA). Cells were maintained in Dulbecco’s modified Eagle’s medium supplemented with 10% (A549, HEK293T, EA.hy926, Gibco) or 5% (Vero, Atlas Biologicals) fetal bovine serum (FBS). Where necessary, cells were selected with puromycin (2 µg/mL) or blasticidin (8 µg/mL). Key small molecule inhibitors used in this study were: NIM811 (MedChemExpress HY-P0025), cyclosporin A (Tocris Bioscience #1101), FK506 (MedChemExpress HY-13756), and juglone (MedChemExpress HY-N6949).

*Ixodes scapularis*-derived embryonic (ISE6) cells were a gift from Uli Munderloh (University of Minnesota, MN, USA). Cells were cultured in L15C300 medium supplemented with 10% heat-inactivated FBS, 10% tryptose phosphate broth, and 0.1% bovine lipoprotein concentrate. Cells were maintained in non-vented T25 flasks at 34°C.

### Propagation and transformation of *R. parkeri*

All strains used in this study are wild-type (WT) or transformed derivatives of *R. parkeri* Portsmouth generously provided by Christopher Paddock (Centers for Disease Control, USA). General procedures for routine propagation of *R. parkeri* have been described previously^11^. Briefly, *R. parkeri* stocks were expanded for use by rocking infection of Vero cell monolayers in T25-T175 flasks, followed by incubation for ∼72 h at 37°C in DMEM + 2% FBS until the host cell monolayer was >90% rounded by light microscopy. *R. parkeri* were then isolated by bead beating, further purified by passage through a 2 µm glass microfiber filter (Cytiva), and stocks were resuspended in brain-heart infusion (BHI) media (BD Difco) and stored as 50-300 µL aliquots at -80°C. Aliquots were titered by plaque assays as previously described^49^, and limited to a maximum of six freeze-thaw cycles for use in experimental infections.

For transformation of *R. parkeri*, bacterial were freshly isolated from an infected T25 flask (for pRAM18dRA (empty vector) and pRAM18dRGA (one copy of GFPuv)) or T75 flask (pEG03 (one copy of AausFP1^87^), gift of Erin Goley (Johns Hopkins University, USA)) and electroporated with 1 µg or 6 µg, respectively, of the indicated plasmids as previously described^10^. Electroporated bacteria were recovered by immediate infection of fresh Vero monolayers and monitoring for plaque formation under rifampicin (200 ng/mL) selection. Bacteria from successfully recovered wells were plaque-purified as previously described to isolate clonal transformants for expansion. *R. parkeri* expressing TagBFP or tandem copies of GFPuv and the isolation of *sca2*::Tn *R. parkeri* transposon mutant have been previously reported^11,29^.

### Molecular cloning

A list of vectors used in this study is provided in **Table S5**. All vectors were sequenced-confirmed by whole-plasmid sequencing (Plasmidsaurus).

For pRDA_550 knockout vectors, synthesized gRNA cassettes (Millipore) harboring tandem gRNAs against each gene of insert from the Humagne C+D gRNA library were inserted into BsmBI-digested pRDA_550 by standard Golden Gate cloning procedures. Guide sequences used in this study are listed in **Table S6**.

For pFUW2IB-based lentiviral rescue constructs, an ORF containing *H. sapiens* PPIA C-terminally fused to dsRed-Express2 with the previously reported linker GSGGSGGSGGQSTVPRARDPPVAT^32^ was synthesized (Twist Biosciences) and cloned into NheI/EcoRI-digested pFUW2IB by standard restriction cloning. The PPIA^R55A^-dRE2 construct was constructed by round-the-horn PCR followed by DpnI digestion, ligation and transformation into DH5α *E. coli*.

For *R. parkeri* Sca2 protein purification constructs, *sca2* regions were amplified from *R. parkeri* Portsmouth gDNA and cloned by Gibson assembly into the pGEX-6P3 backbone. Truncations and point mutations, including the pET3A-OSF-PPIA(R55A) construct, were constructed by round-the-horn PCR followed by DpnI digestion, PNK phosphorylation, ligation, and transformation into DH5α *E. coli*.

### Flow cytometry of *R. parkeri*-infected cells

A549 parental or knockout cells were seeded in 12 well plates at 2×10^5^ cells/well 2 days before infection. On the day of infection, cells were spinfected with the indicated strains of *R. parkeri* at an MOI ∼3 (0.1-10 for MOI titration experiments) at 200 rcf for 5 min at RT. Cells were incubated at 33°C to allow infection to proceed. At the indicated timepoints, cells were trypsinized and pelleted in 1.5 mL tubes at 1200 rcf for 5 min at RT. Media was aspirated from pelleted cells, which were then resuspended in 500 µL fixative (4% PFA in PBS or 100% ice-cold methanol), vortexed briefly, and incubated on ice for 15 min. For methanol fixation, tubes were first briefly dragged along a rack to avoid subsequent clumping upon methanol addition. After fixation, 500 µL sterile FACS buffer (1xPBS + 1% BSA + 0.02% sodium azide) was added, and cells were pelleted at 300 rcf for 5 min. Cells were washed once with FACS buffer and transferred to filter cap round bottom tubes (BD) for flow cytometry.

For antibody staining of PFA-fixed infected cells, after washing with FACS buffer, cells were re-pelleted and permeabilized with 0.5% Triton-X 100 in FACS buffer on ice for 10 min. Permeabilized cells were washed with FACS buffer and then stained with mouse 14-13 anti-*Rickettsia* (gift of Ted Hackstadt) diluted in FACS buffer on ice for 30 min. Cells were washed twice with FACS buffer and then stained with goat anti-mouse Alexa Fluor 647 (Invitrogen #A-21236) diluted in FACS buffer on ice for 30 min. Cells were washed twice with FACS buffer and transferred to a filter cap 5 mL round bottom tube for flow cytometry. All flow cytometry analysis was performed on a FACSCanto II instrument (BD). For all GFP derivatives, a 488 nm laser was used with an LP 502 and BP 530/30 configuration. For TagBFP, a 405 nm laser was used with a BP 450/50 configuration. For Alexa Fluor 647, a 633 nm laser was used with a BP 660/20 configuration. All samples were gated for acquisition on single cells using SSC-A, FSC-A and FSC-H. For gating fluorescence, samples were compared to uninfected control wells from the same plate. Data analysis was performed with FlowJo 10 (BD).

### Genome-wide CRISPR/Cas12a screen and data analysis

The A549-enAsCas12a stable cell line was generated by transducing parental A549 cells with pRDA_174 lentivirus, followed by blasticidin selection for 14 d. enAsCas12a expression was verified by Western blot against the HA epitope tag on enAsCas12a and activity was verified by subsequent transduction with pRDA_221 harboring a GFP expression cassette and a GFP-targeting gRNA, followed by assessment of GFP fluorescence reduction relative to parental A549 cells by flow cytometry. To generate the knockout library, 9×10^7^ A549-enAsCas12a cells were transduced with lentivirus carrying the Humagne C+D (CP1882) human genome-wide knockout library (500x representation at 30% predicted transduction efficiency). The calculated transduction efficiency was 35%. Transduced cells were selected with puromycin for 7 d, maintaining a minimum population size of 6×10^7^ to avoid bottlenecks. After 1 week, the minimum population size was lowered to 2×10^7^ cells. For the first replicate of the screen, 10 T175 flasks of the library were seeded at 6×10^6^ cells in 35 mL total media per flask. 72 h after seeding, flasks were infected with *Rp*AausFP1. At 48 hpi, infected cells were harvested by trypsinization followed by centrifugation at 1200 rcf for 5 min. Cell pellets were pooled and fixed by resuspension in 100% ice-cold methanol by drop-by-drop addition with continuous vortexing. Resuspended cells were incubated on ice for 15 min and then washed in FACS buffer. Cells were diluted to 1×10^7^/mL in FACS buffer and sorted on a FACSAria II (BD) to isolate the top and bottom 10% GFP fluorescence bins relative to a fixed, uninfected sample of the library. At least 6×10^6^ cells were isolated in each bin and used for gDNA isolation (Machery-Nagel Nucleospin) alongside an equivalent sample of unsorted cells. All isolated gDNA (>10 µg/sample) was used for PCR amplification of the gRNA cassette and sequencing, performed by the Broad Institute Genetic Perturbation Platform. The second replicate of the screen was performed with a later passage of the same library. The screen was conducted identically to the first replicate, with the only modification being a higher cell concentration (1.5×10^7^/mL) used for FACS. Screen analysis was performed with Apron (Broad Institute) (see “Bioinformatic Analyses” section for further details).

### Generation of knockout cell line pools, clones and rescue lines

For all knockout lines, the enAsCas12a-based all-in-one lentiviral knockout vector pRDA_550 was used. The lentiviral expression vector pFUW2IB was used to express specific constructs under the control of the UbC promoter. To generate lentivirus, HEK293Ts were transfected with 280 ng pRDA_550 or pFUW2IB vector and 140 ng each of the pMDLg/pRRE, pRSV-REV and pCMV-VSV-G packaging vectors with LT-1 transfection reagent (Mirus Bio #2304) according to standard protocols and lentivirus was collected in media supplemented with 1% BSA. Lentiviral supernatants were collected 48 h post-transfection and filtered through 0.45µm PES filters (Genesee Scientific #25-246) before transduction. A549s transduced with the indicated pRDA_052 derivative harboring a gRNA from the Humagne C+D library were selected with puromycin for at least 7 d before knockout validation (via immunoblot) and/or phenotyping. For single-cell cloning of *PPIA*ko lines, clones were isolated by limiting dilution in 96-well plates followed by growth for 14 d in media containing puromycin. Wells containing single colonies were trypsinized and transferred to larger culture plates for phenotyping by immunoblot for PPIA. Clones with obviously altered cell morphology or growth kinetics were not propagated. To generate *PPIA*ko rescue lines or parental A549 lines expressing dRE2-tagged proteins, cells were transduced with pFUW2IB derivatives and selected with blasticidin for 14 d. dRE2-tagged protein-expressing lines were single-cell cloned as described for the *PPIA*ko clones. For *PPIA*ko rescue lines, lentiviral *PPIA* ORFs were altered with at least 3 silent mutations within the first 10 nt of the seed sequence to render them enAsCas12a-resistant (listed in **Table S6**).

### General confocal immunofluorescence (IF) microscopy

All IF assays used the same standard fixation and staining protocol, with variations noted below for each specific assay. Host cells were seeded in 24-well tissue culture-treated plastic plates (Genclone) on sterile glass #1.5 coverslips (Fisherbrand) and allowed to attach for at least 24 h. At the indicated infection endpoint, cells were fixed with 4% PFA for 15 min at RT and then washed three times with 1xPBS. Fixed coverslips were quenched with 0.1 mM glycine for 10 min and then washed three times with 1xPBS. Cells were then permeabilized with 0.5% Triton-X 100 in PBS for 5 min, after which cells were washed with 1xPBS and blocked for 30 min with 2% BSA/PBS. Blocking solution was then replaced with primary antibodies diluted in 2% BSA/PBS for 1 h at RT. Cells were washed by soaking three times for 5 min with 1xPBS and secondary antibodies diluted in 2% BSA/PBS added for 1 h at RT. Coverslips were washed and mounted on glass slides using ProLong Gold antifade mountant (Invitrogen). Mounted slides were dried overnight in the dark and sealed with nail polish for imaging. Images were acquired on an Olympus IXplore Spin microscope system with a Yokogawa CSU-W1 spinning disk unit and an ORCA-Flash4.0 sCMOS camera using either 60x UPlanSApo (1.30 NA) (spread assays) or 100x UPlanSApo (1.35 NA) (all other assays) objectives. At least 10 fields of view per coverslip were acquired by scanning the *R. parkeri* channel to image solely based on bacterial density. Image analysis was performed in ImageJ.

### Infectious focus (spread) assay

A549 cells were seeded at 2×10^5^ cells/well in 24 well plates 2 d prior to infection. Cells were spinfected with MOI ∼0.001 WT *R. parkeri* and incubated at 33°C for 1 h, after which media was aspirated, wells washed three times with sterile 1xPBS, and fresh media containing 10 µg/mL gentamicin was added to each well. Plates were incubated for an additional 27 h at 33°C before fixation. Primary antibodies used were: rabbit chimeric 14-13 anti-*Rickettsia*^88^ (gift of Ted Hackstadt (Rocky Mountain Laboratories/NIAID, USA)), mouse anti-β-catenin (CST #2677S). Secondary antibodies and stains used were: Hoescht (Invitrogen H3570), goat anti-rabbit Alexa Fluor 488 (Invitrogen A-11001), and goat-anti mouse Fcγ Alexa Fluor 568 (Jackson ImmunoResearch #115-605-071). At least 20 infection foci per coverslip were imaged for quantification.

### Actin tail assay

A549 cells were seeded at 2×10^5^ cells/well in 24 well plates 2 d prior to infection. Cells were spinfected with MOI ∼0.5 WT *R. parkeri* and incubated at 33°C for 48 h before PFA fixation as described above. Cells were stained as for the infectious focus assay with the addition of phalloidin Alexa Fluor 647 (Invitrogen #A22287). Tails were counted by first segmenting single *R. parkeri* with MicrobeJ as described below, then scoring for actin tails. Tails were only counted if they exceeded the length of the associated bacterium.

### Drug treatment and washout assays

For single timepoint drug treatments, A549 cells were seeded at 2×10^5^ cells/well in 24-well plates 2 d prior to infection. Cells were spinfected with MOI ∼0.5 WT *R. parkeri* and incubated at 33°C for 24 h, at which point media was removed, cells washed once in 1xPBS and fresh media containing the indicated drugs at the indicated concentrations was added. Cells were returned to 33C and incubated for an additional 24 h. Cells were fixed at 48 hpi. Assays assessing the effect of drugs on actin tails were stained identically to the actin tail assay. Assays assessing the effect of drugs on Sca2 surface exposure used the following primary antibodies: mouse 14-13 anti-*Rickettsia* and rabbit anti-*R. parkeri* Sca2 (in-house). Secondary antibodies and stains used were: Hoescht, goat anti-mouse Alexa Fluor 488, goat anti-rabbit Alexa Fluor 568 (Invitrogen #A-11004), and phalloidin Alexa Fluor 647. For CsA washouts, at 1 hpi cells were washed with 1xPBS and then media containing 10 µM CsA was added. At varying washout timepoints, media was removed, cells were washed three times with 1xPBS, and CsA-free media was added. At 48 hpi, cells were fixed and stained as described above.

For drug treatment of ISE6 cells, cells were seeded at a density of 5×10^5^ cells/well in 24-well plates on glass coverslips 4 d prior to infection. Cells were spinfected (300 rcf, 5min) with *R. parkeri* at an MOI of 1. Plates were placed in airtight containers and incubated at 34°C for 1 h, before being gently washed with 1X PBS and changed into fresh L15C300 media containing 5 µM CsA, 5 µM NIM811 or DMSO vehicle. Cells were incubated for 48 h in airtight containers at 34°C before being fixed with 4% PFA at 48 hpi. Cells were stained as for the actin tail assay.

### PPIA-dRE2 localization assays

A549 cells were seeded at 4×10^4^ cells/well in 24-well plates 2 d prior to infection. Cells were spinfected with MOI ∼2.5 WT *R. parkeri* and incubated at 33°C. For treatment with CsA, 10 µM CsA or DMSO was added to cells as described above at 24 hpi. At 48 hpi, cells were fixed with 4% PFA for 1 h in the dark at RT before washing and staining by the standard protocol described above. The following primary antibodies were used: mouse 14-13 anti-*Rickettsia* and rabbit anti-*R. parkeri* Sca2. The following secondary antibodies were used: goat anti-mouse Alexa Fluor 405 and phalloidin Alexa Fluor 488.

### Lysozyme permeabilization immunofluorescence assay for Sca2

The differential permeabilization assay was performed as previously described^49^. Briefly, A549 cells were seeded, infected and fixed as described for the actin tail assay with duplicate coverslips seeded per condition for staining with and without lysozyme permeabilization. All coverslips were first stained with the standard IF protocol using the primary antibody mouse 14-13 anti-*Rickettsia* and the secondary antibody goat anti-mouse Alexa Fluor 488, Hoechst, and phalloidin 647. For non-permeabilized coverslips, this also included the rabbit anti-*R. parkeri* Sca2 primary and goat anti-rabbit Alexa Fluor 568 antibodies, after which staining was complete. For permeabilized coverslips, after the first round of staining, samples were re-fixed with 4% PFA for 5 min at RT and stored in 1xPBS overnight at 4°C. Samples were re-quenched with 100 mM glycine as described above, then permeabilized by incubation in lysozyme reaction buffer (0.8×PBS, 50 mM glucose, 5 mM EDTA, 0.1% Triton X-100, 5mg/mL lysozyme (Sigma #L6876)) for 20 m at 37°C. Permeabilized samples were washed three times with 1xPBS and then stained as described above with the rabbit anti-*R. parkeri* Sca2 primary and goat anti-rabbit Alexa Fluor 568 secondary antibodies. Image analysis was performed with MicrobeJ^89^ version 5.13p(1). Single bacteria were detected using the 14-13 anti-*Rickettsia* signal with custom settings: smoothed morphology, area 0.4-3.5, length 0.5-3.5, width 0.4-1, circularity 0.5-1, angularity 0-0.5 (all µm). Clumps of bacteria, bacteria in the nucleus (which have abnormally high rates of actin-based motility^90^), as well as any cells along the edge of the image were excluded. Segmented bacterial outlines were used to manually score Sca2-positive *R. parkeri*. For measuring total levels of Sca2 in permeabilized bacteria, MicrobeJ was used to calculate the sum intensity of the Sca2 channel in segmented *R. parkeri*. This total was then normalized by bacterial area.

### Immunoblotting

Uninfected samples for verifying PPIA knockout were harvested during routine passaging. For samples from infected cells, A549s were at 2×10^5^/well in 12 well plates 2 d prior to infection. Cells were infected with WT or *sca2*::Tn *R. parkeri* at MOI ∼3-5 by spinfection and incubated at 33°C until harvesting at 48 hpi. Samples from uninfected or infected plates were processed identically for Western blotting. Cells were trypsinized, pelleted at 1200 rcf for 5 min at RT and resuspended in 1x SDS loading buffer (150mM Tris-HCl pH 6.8, 6% SDS, 0.3% bromophenol blue, 30% glycerol). Samples were boiled for 10 min and electrophoresed on 12% SDS-PAGE gels. Proteins were transferred to PVDF membranes using semi-dry transfer (Bio-Rad Trans-Blot) using the preset 1.5 mm gel protocol. Membranes were blocked in 5% milk in TBST (1x TBS with 0.1% Tween-20) for 1 h at RT and then primary antibodies were added for overnight incubation at 4°C. Membranes were washed three times for 10 min each with TBST, and secondary antibodies were added for 1 h at RT followed by three additional 10 min washes with TBST. Membranes were incubated with chemiluminescent substrate (Thermo # A43840) for imaging. Primary antibodies used were: rabbit anti-PPIA (Thermo #PA1-025), mouse anti-alpha tubulin (Sigma #T6199), rabbit anti-Sca2 (in-house purified from polyclonal serum), mouse anti-RpoB (BioLegend #663905). Secondary antibodies used were HRP-conjugated goat anti-mouse or rabbit IgG (Jackson Immunoresearch #115-035-003 (mouse) and #111-035-003 (rabbit)). All antibodies were diluted in 2.5% milk in TBST.

### Purification of GST-tagged *R. parkeri* Sca2

To purify recombinant GST-Sca2 proteins, overnight cultures of Rosetta2 *E. coli* transformed with pGEX-6P3-Sca2 derivatives were diluted 1:500 into 1 L cultures of 2xYT media with ampicillin and chloramphenicol. Cultures were grown at 37°C at 200 rpm until OD ∼0.8, at which point they were induced with 0.1mM IPTG and grown at 18°C overnight. Cultures were harvested by centrifugation at 5000 rcf for 15 min and resuspended in 10 mL/L lysis buffer (50 mM Tris pH 8, 150 mM KCl, 0.1% Tween-20) freshly supplemented with 1 mM PMSF, 20 µg/mL Dnase I and protease inhibitors (Roche cOmplete Mini). Cells were incubated with 1 mg/mL lysozyme for 15 min at 4°C. Then, 1 mM DTT was added, and cells were lysed by sonication followed by centrifugation at 30,000 rcf for 30 min. Clarified lysate was rocked overnight at 4°C with glutathione 4B Sepharose (Cytiva) before gravity column purification. Resin was washed with 2 column volumes of high-salt wash buffer (50 mM Tris pH 8, 500 mM KCl, 1 mM DTT, 1 mM PMSF) followed by 6 column volumes of standard wash buffer (50 mM Tris pH 8, 150 mM KCl, 1 mM DTT, 1 mM PMSF) and elution in freshly prepared GST elution buffer (50 mM Tris pH 8, 150 mM KCl, 1 mM DTT, 10 mM reduced glutathione). Fractions were pooled, concentrated by centrifugation and further purified by size exclusion chromatography on a Superose 6 10/300 GL column (Cytiva) into the final buffer (50 mM Tris pH 8, 150 mM KCl, 1 mM TCEP). Pure protein fractions were pooled, re-concentrated and stored at -80°C for use. GST did not require purification by size exclusion and was purified by an identical glutathione affinity procedure using buffers containing NaCl rather than KCl and omitting the high-salt wash step. All purification steps were performed at 4°C.

### Purification of OSF-tagged human PPIA

One-STrEP-FLAG (OSF)-tagged CypA (PPIA) was purified largely as previously described^91^. Overnight cultures of BL21(DE3) *E. coli* transformed with pET3a-OSF-CypA were diluted 1:1000 into 1 L cultures of ZYP-5052 autoinduction media (10 g/L tryptone, 5 g/L yeast extract, 1×5052 (0.05% d-glucose, 0.5% glucose, 0.2% α-lactose), 1xNPS (25 mM (NH_4_)_2_SO_4_, 50 mM KH_2_PO_4_, 50 mM Na_2_HPO_4_)). Cultures were grown at 37°C, shaking at 200 rpm overnight. Cultures were harvested by centrifugation at 5000 rcf for 15 min. and resuspended in 10 mL/L culture lysis buffer (50 mM Tris pH 8, 50 mM NaCl, 0.2% deoxycholate) freshly supplemented with 2.5 nmol avidin, 10 mM β-mercaptoethanol (β-ME), 1 mM PMSF, 20 µg/mL Dnase I, and protease inhibitors (Roche cOmplete Mini). Cells were lysed by sonication and lysates were clarified by centrifugation at 30,000 rcf for 30 min. Clarified lysates were incubated with Streptactin XT 4flow resin (IBA Lifesciences) for 1 hr at 4°C with rotation before gravity column purification. Isolated resin was washed with 6 column volumes of wash buffer (100 mM Tris pH 8, 150 mM NaCl, 10 mM β-ME) before elution with elution buffer (wash buffer + 2.5mM d-desthiobiotin). Fractions were pooled, concentrated and further purified by size exclusion chromatography using a Superdex 200 Increase 10/300 GL column (Cytiva). Pure protein fractions were pooled, re-concentrated and stored at -80°C. All centrifugation steps were performed at 4°C.

### *In vitro* pulldown assays

GST pulldowns were adapted from a previously published protocol^92^. The GST-Sca2 size exclusion buffer was used as the buffer for all binding assays. 4-10 µM of GST or GST-Sca2 and OSF-PPIA were mixed in equal ratios in a total volume of 50-100 µL, depending on the scale of the assay. Protein mixtures were incubated on ice for 15 min, after which 50 µL of equilibrated glutathione 4B Sepharose (Cytiva) was added to the mixture and incubated for a further 10 min. Bound protein on beads was then collected by centrifugation at 300 rcf for 5 min, followed by three washes with 10x bead volume of binding buffer. Proteins were eluted by incubation with GST elution buffer (50 mM Tris pH 8, 10 mM reduced glutathione) for 5 min on ice. Samples were mixed with 3x SDS loading dye and analyzed by SDS-PAGE followed by staining with SimplyBlue Safestain (Life Technologies). For reactions where CsA was added, the drug or DMSO vehicle was added at the beginning of the first incubation.

### In vivo R. parkeri mouse infections

All mouse experiments were conducted at UC Irvine with the approval of the UCI Institutional Animal Care and Use Committee (IACUC) (Protocol #AUP-25-004). All mice were healthy at the time of infection and provided food and water *ad libitum*. All experiments were performed with WT C57BL/6J mice (Jackson Labs). No mice were exposed to antibiotics at any time. For each experimental group, 4-6 age-matched littermates were randomly assigned to receive either vehicle or NIM811, with similar numbers of males and females in each group. To prepare inocula, WT or *sca2*::Tn *R. parkeri* were prepared by density gradient centrifugation as previously described^53^ and then thawed on ice and pelleted at 10,000 rcf for 2 min at 4°C. Pellets were resuspended in cold sterile PBS to a concentration of 2.5×10^7^ plaque-forming units (PFU) per mL and kept on ice during injections. Mice were exposed to a heat lamp while in their cages for 5 min and then each mouse was moved to a restrainer (Braintree, TB-150 STD). The tail was sterilized with 70% ethanol and 200 μl bacterial suspension (5×10^6^ PFU) per mouse was injected into the lateral tail vein with a 29-gauge needle. For drug treatment, mice were treated with 50 mg/kg NIM811 or vehicle control at -1, 0, 1, and 2 days post-infection (dpi). NIM811 was diluted to a concentration of 125 mg/mL in DMSO and frozen into aliquots to limit freeze-thaw cycles before use. Solubilized NIM811 was thawed at room temperature and diluted with sterile-filtered corn oil (Sigma-Aldrich C8267) to deliver 50 mg/kg (∼100 µL per mouse). The solution was vortexed thoroughly before mice were injected intraperitoneally using 29-gauge needles.

To quantify organ-specific bacterial burdens, mice were euthanized at 3 dpi and doused with ethanol before dissection. Organs were kept on ice in 4 mL (spleen) or 8 mL (liver) of sterile PBS and were homogenized for ∼12 s using an immersion homogenizer at ∼23,000 rpm. Organ homogenates were centrifuged at 260 rcf for 5 min to pellet tissue debris, and then 20 µL aliquots of the supernatant were serially diluted onto confluent Vero monolayers in 12-well plates, and then spinfected at 300 rcf for 5 min at room temperature. At 1 hpi, carbenicillin and amphotericin B were diluted into each well to a final concentration of 50 and 1 μg/mL, respectively. At 18 hpi, media containing antibiotics were aspirated and replaced with 2 mL of overlay solution (DMEM + 5% FBS, 2.4% Avicel, and 1 μg/mL amphotericin B). At 6-8 dpi, monolayers were fixed with 2 ml of 7% PFA per well and incubated for >30 min before staining with 1.25% crystal violet solution for >15 min before washing generously with water to count plaques. PFU/organ was calculated by multiplying the ratio of the total homogenate volume to sample volume.

### Pathogen-specific *in vitro* infections

#### Listeria monocytogenes *and* Shigella flexneri

*L. monocytogenes* 10403S was grown on brain heart infusion (BHI) agar at 30°C for 18-20 hr. The following day, 5 colonies were used to inoculate 2.5 mL BHI broth and incubated as a stationary culture overnight at 30°C for 18-20 hr. 1 mL of overnight culture was pelleted at 16,200 rcf for 1 min and pellets were resuspended in 1 mL PBS. A549 cells seeded at 1.25 x 10^5^ cells/well in 24-well plates 5 d prior were infected at an MOI of 0.5, centrifuged at 300 rcf for 5 min and incubated for 1 hr at 37°C + 5% CO2 to facilitate invasion. Media was then removed and the cells were washed twice with 0.5mL 1x PBS before being given fresh 0.5 mL DMEM + 10% FBS + 10 µg/mL gentamicin along with the indicated concentration of NIM811 or DMSO vehicle control and incubated for the indicated time points for enumeration of colony-forming units (CFUs).

*S. flexneri* 2457T was grown on tryptic soy agar + 0.01% Congo Red at 37°C for 18 hr. A single red colony was used to inoculate 4 mL tryptic soy broth and incubated at 37°C while shaking for 18 hr. The following day, cultures were diluted 1:200 and grown at 37°C while shaking until OD_600_ = 0.4 – 0.6 (∼2.5 hr). 1 mL of culture was pelleted at 16,200 rcf for 1 min, and pellets were resuspended in 1 mL PBS. A549 cells were then infected as for *L. monocytogenes*, with the only modification being an MOI of 1 and the use of 25 µg/mL gentamicin in the post-wash media. To enumerate CFUs, cells were washed twice with 0.5 mL 1xPBS and lysed with 300 µL 0.1% TritonX-100 in PBS for 5 min. Lysates were diluted by 10-fold serial dilutions, and 10 µL of each dilution was drip plated on each bacterium’s respective media and grown at 37°C overnight. Dilutions that contained between 2 – 100 colonies were counted and used to calculate CFU/well. At least two technical replicate wells were included per condition.

#### Chlamydia trachomatis

HeLa cells were grown at 37°C with 5% CO_2_ in RPMI 1640 medium (ThermoFisher Scientific) supplemented with 10% FBS (Gibco) and infected with *C. trachomatis* serovar L2 (LGV 434/Bu) largely as previously described^93,94^. HeLa cells were infected at an MOI of 1 on ice and after 30 min, the inoculum was removed and replaced with fresh RPMI. At 5 hpi, cells were treated with DMSO (vehicle) or 10 µM NIM811. To determine infectious bacterial burdens at 48 hpi, host cells were lysed in water, serially diluted, and applied to fresh HeLa cell monolayers to determine the number of infectious forming units. Each independent experiment was performed with three technical replicate wells.

#### Orientia tsutsugamushi

For *O. tsutsugamushi* work, uninfected HeLa human cervical epithelial cells (CCL-2; ATCC) and EA.hy926 human umbilical vein cells (CRL-2922; ATCC) were maintained as described previously^95^. *O. tsutsugamushi* strain Ikeda was maintained in HeLa cells^95,96^ and host cell-free bacteria were isolated for infection studies in EA.hy926 cells as described^97^. All experiments were performed at an MOI of 5-10 resulting in at least 90% host cell infection, confirmed by immunofluorescence microscopy as described previously^97^. For infections, EA.hy926 cells were seeded in 24-well plates to achieve a confluency of 100%. To synchronously infect cells, *O. tsutsugamushi*-infected EA.hy926 cells were centrifuged at 1000 rcf for 3 min and incubated at 35°C in 5% CO_2_ for 1 hr before inocula were replaced with fresh complete DMEM containing l-glutamine, 4.5 g d-glucose and 100 mg sodium pyruvate (Gibco) supplemented with 10% (vol/vol) heat-inactivated fetal bovine serum (FBS) (Gemini Bioproducts), 1X minimal essential medium containing nonessential amino acids (Gibco) and 15 mM HEPES (Gibco). At 4 hpi, medium was replaced with complete DMEM plus 10 µM NIM811 and incubated for 72 h. Lysates for GE of host cell-free *O. tsutsugamushi* inocula and of infected EA.hy926 cells at 1 and 72 hpi were collected as described previously^97,98^. Briefly, samples were dislodged by trypsin, diluted in H_2_O, incubated at 100°C for 15 min, and stored at -20°C until analysis. GE was quantified by qPCR analysis using primers to detect *O. tsutsugamushi* strain Ikeda *tsa56* (OTT_RS04590)^96^ with PerfeCTa SYBR Green Fastmix (Quantbio) and a CFX384 Real Time PCR Detection System (Bio-Rad). Thermal cycling conditions were 95°C for 30 s followed by 40 cycles of 95°C for 10 s to 54°C for 10 s, and a 65-95°C melt curve. *O. tsutsugamushi* yield was calculated by subtracting the GE/mL at 72 hpi by the GE/mL of the *O. tsutsugamushi* inoculum for the corresponding experiment. Experiments were performed in triplicate with three technical replicates.

### Bioinformatic analyses

CRISPR screen analysis was performed with Apron (beta version, Broad Institute). Apron uses aggregated log-normalized read constructs for guides targeting each gene to calculate fold changes based on comparison to a reference distribution. P-values were calculated from the standard distribution of gene z-score deviation from fold changes of non-target and intergenic-targeting gRNAs. False discovery rates were calculated using the Benjamini-Hochberg procedure on p-values within each condition. Two analyses were performed. The first compared the bulk output libraries (pooled reads from low, high and unsorted bins) in each replicate to the input plasmid DNA to verify library complexity and essential gene dropout. The second compared the high to low fluorescence bins in each replicate to identify hits (see **Fig. S2**).

For STRING network analysis, the top 100 and bottom 100 hits were combined and used to generate a functional and physical interaction network. Clusters were identified with DBSCAN clustering using an epsilon parameter of 5. Structural prediction was performed with the AlphaFold3 server (alphafoldserver.com)^99^. The highest-ranked structure was used for visualization with ChimeraX^100^. PAE plots were generated with PAE Viewer^101^. Assessment of proline isomerase conservation was performed with manual Annotree^102^ (version r214 using GTDB R214 database) lookups of PF00160, PF00254 and PF13145 at an e-value less than 0.00001. A cladogram for visualization was generated with the BV-BRC pathogen phylogenetic tree tool^103^ using 72 conserved genes and the RAxML phylogenetic tree algorithm. Specific strains used for the cladogram are listed in **Table S3**. Annotree absences were double-checked with literature searches to identify any annotation gaps. Species lacking both an Annotree annotation and any published reports were classified as “absent” for the indicated isomerase family.

### Statistics and replicates

Replicate information and specific statistical tests are indicated on each figure legend. Biological replicates for tissue culture infections were defined as wells that were infected and harvested or fixed on different days. Unless otherwise stated, graphed summary values represent replicate means with standard deviation error bars. Unless otherwise stated, p-values greater than 0.05 were considered not statistically significant. Statistical analyses are outlined in each figure and were performed with Microsoft Excel or Graphpad Prism.

## Acknowledgements

We thank members of the Lamason Lab for helpful discussions throughout this project and Gregory Babunovic, Karthik Hullahalli, and Alyson Warr-Manteiga for critical reading of the manuscript. We also thank Seychelle Vos for access to FPLC equipment and Roberto Vásquez-Núñez for technical assistance with protein purification. We also thank the Whitehead Institute Flow Cytometry Core, especially Patrick Autissier and Aditya Rathee, for technical assistance with cell sorting and analysis, and the Broad Institute Genetic Perturbation Platform for technical assistance with library preparation and sequencing.

## Author Contributions

BS and RLL conceived the study. JGD contributed to genetic screen design and feasibility studies. BS conducted the screen, all protein purification work and bioinformatics, and most *R. parkeri* experiments with assistance from CYZ and LEB. AGS performed PPIA localization studies. AAG and TPB carried out *in vivo R. parkeri* infections. PJW carried out *L. monocytogenes* and *S. flexneri* studies. PNM, JK-Y and MMW carried out *C. trachomatis* studies. PEA and JAC carried out *O. tsutsugamushi* studies. LEB additionally performed tick cell infections. BS and RLL wrote the manuscript with review from all other authors.

## Supplementary Tables

**Table S1. Genome-scale CRISPR/Cas12a *R. parkeri* infection screen results.**

**Table S2. DBSCAN cluster analysis of CRISPR screen hits.**

**Table S3: Species and strains used for cladogram generation.**

**Table S4: Annotree results for PF00160, PF00254 and PF13145.**

**Table S5: Bacteria and plasmids used in this study.**

**Table S6: Guide and ORF sequences used in this study.**

## Supplementary Figure Legends

**Figure S1.**
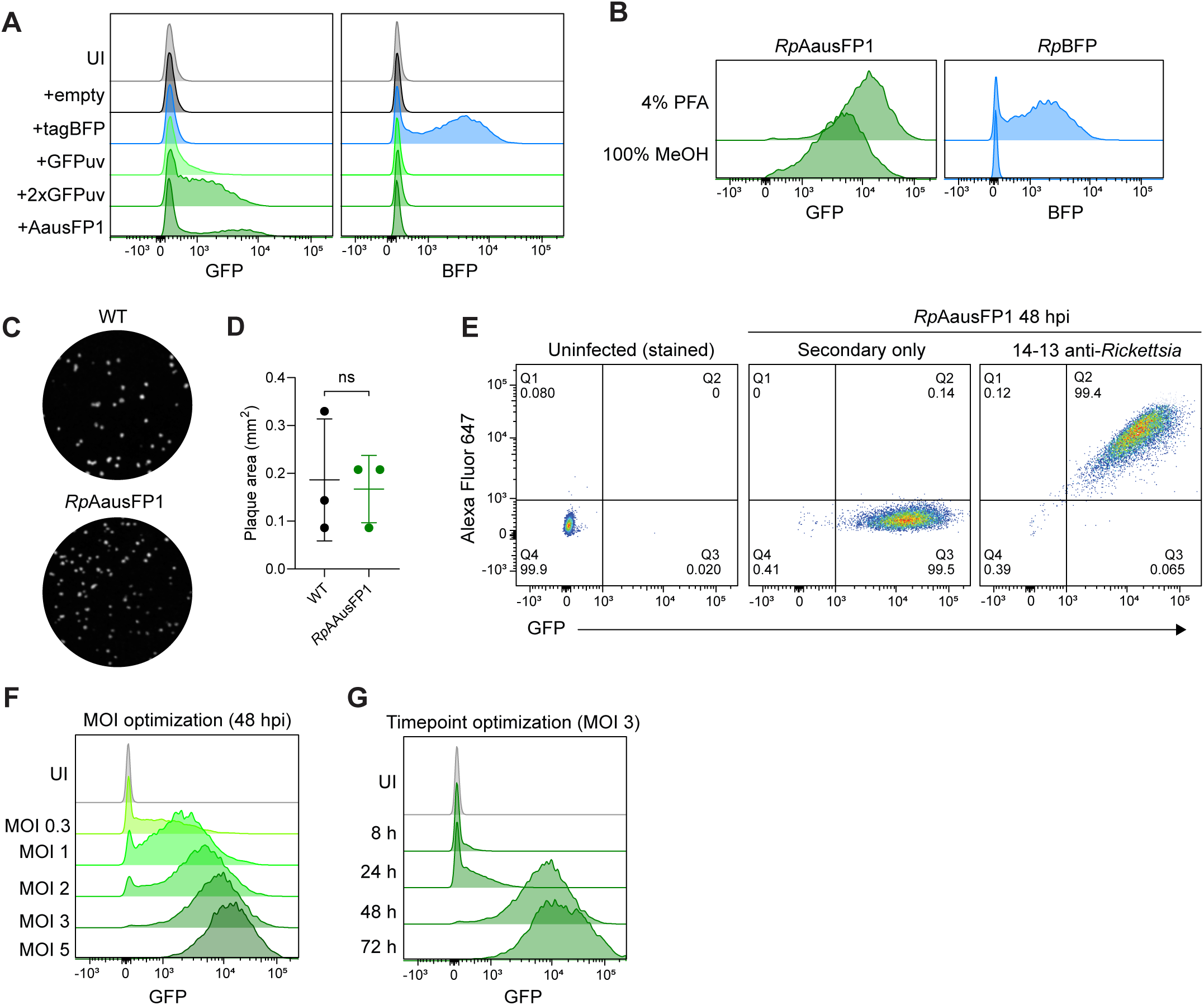
Screen optimization and execution. (**A**): Flow cytometry of A549 cells infected with *R. parkeri* transformed with various fluorescent protein expression constructs at 24 hpi. UI, uninfected. (**B**): Fixation sensitivity of *Rp*AausFP1 and *Rp*BFP strains. (**C** and **D**): Representative plaques (**C**) and quantification (**D**) of Vero cells infected with WT *R. parkeri* or *Rp*AausFP1. Each point represents the mean of >100 quantified plaques from a biological replicate. Means were used to calculate the p-value (Welch’s t-test). (**E**): Correlation of antibody staining and endogenous fluorescence signal in *Rp*AausFP1-infected A549 cells. Samples were stained with a secondary antibody conjugated to Alexa Fluor 647. Stained uninfected cells were used to set gates and a secondary antibody-only infected sample was used as a control for nonspecific binding. (**F**): Optimization of MOI for infections. Only the 48 hpi timepoint is shown. (**G**): Optimization of timepoint for infections. Only the MOI 3 infections are shown (48 hpi curve is the same as in panel F). Y-axes in panels A, B, F and G are all normalized to mode. All flow cytometry plots are representative of at least n=3 biological replicates.

**Figure S2.**
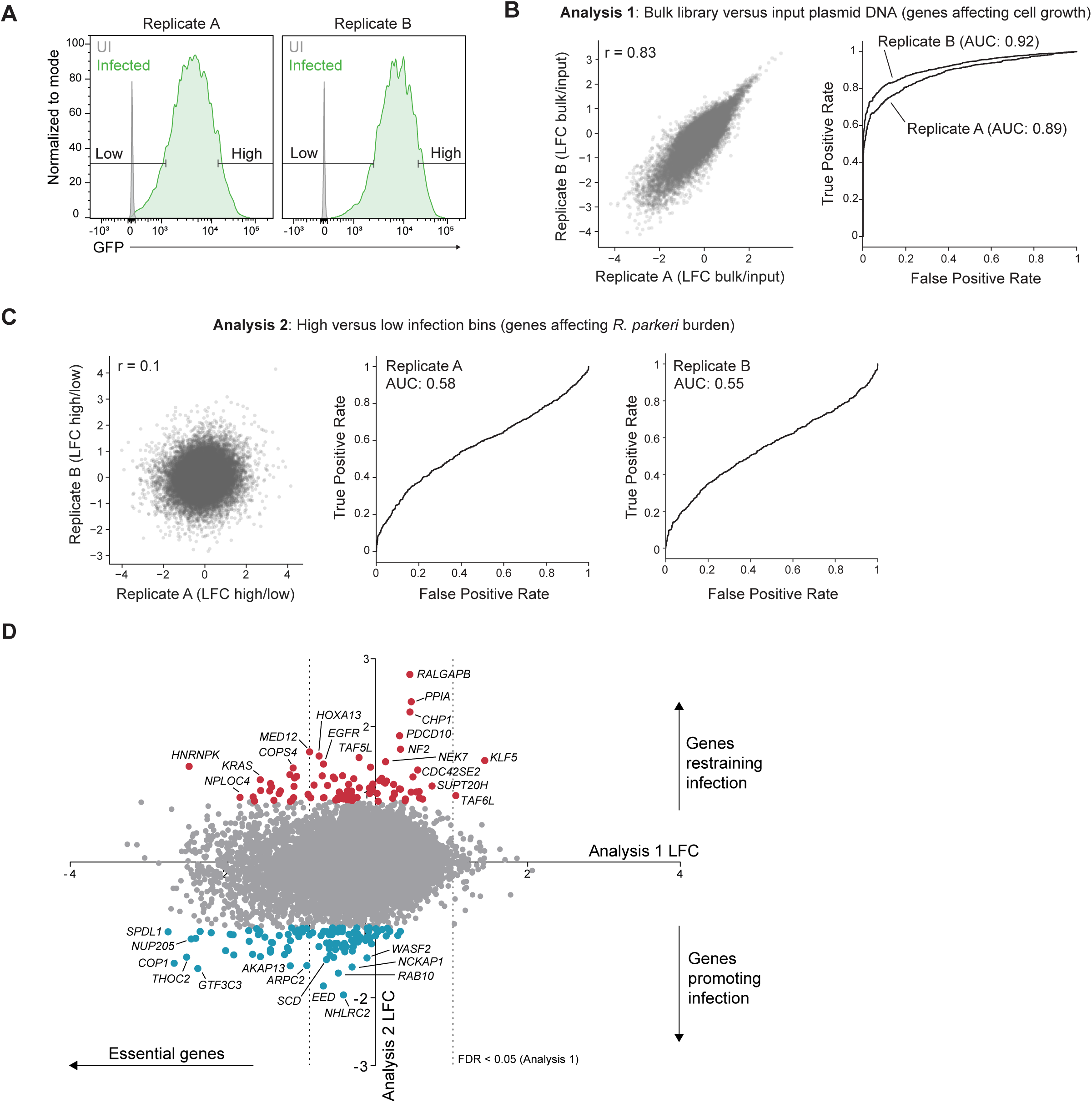
Infection screen results and analysis. (**A**): Flow cytometry of each screen replicate. UI, uninfected. (**B** and **C**): Analysis of screen data comparing either the bulk output libraries versus input pDNA (**B, Analysis 1**) or high versus low bins (**C, Analysis 2**). Left - gRNA-level log2 fold change (LFC) correlation between replicates. Right - gene-level receiver-operator curve(s) for recovery of known essential genes in each replicate. Area under the curves (AUCs) are listed next to each condition. AUC values >0.8 indicate strong separation of positive from negative controls, in this case separation of known essential from known non-essential genes. AUCs are low in panel C because there is no selective pressure against essential genes. r-values shown are Pearson’s correlation coefficient. The r-value is low in panel C because there is no selective pressure for or against most of the library, so most gRNAs behave stochastically. (**D**): Scatter plot of gene LFCs between Analysis 1 and 2. Red and blue points identify genes with FDR < 0.05 (same as Figure 1C). Dashed lines indicate LFC thresholds with FDR < 0.05 for Analysis 1. Genes with strong LFC values for Analysis 1 were generally not selected for validation because they likely have effects on host cell viability.

**Figure S3.**
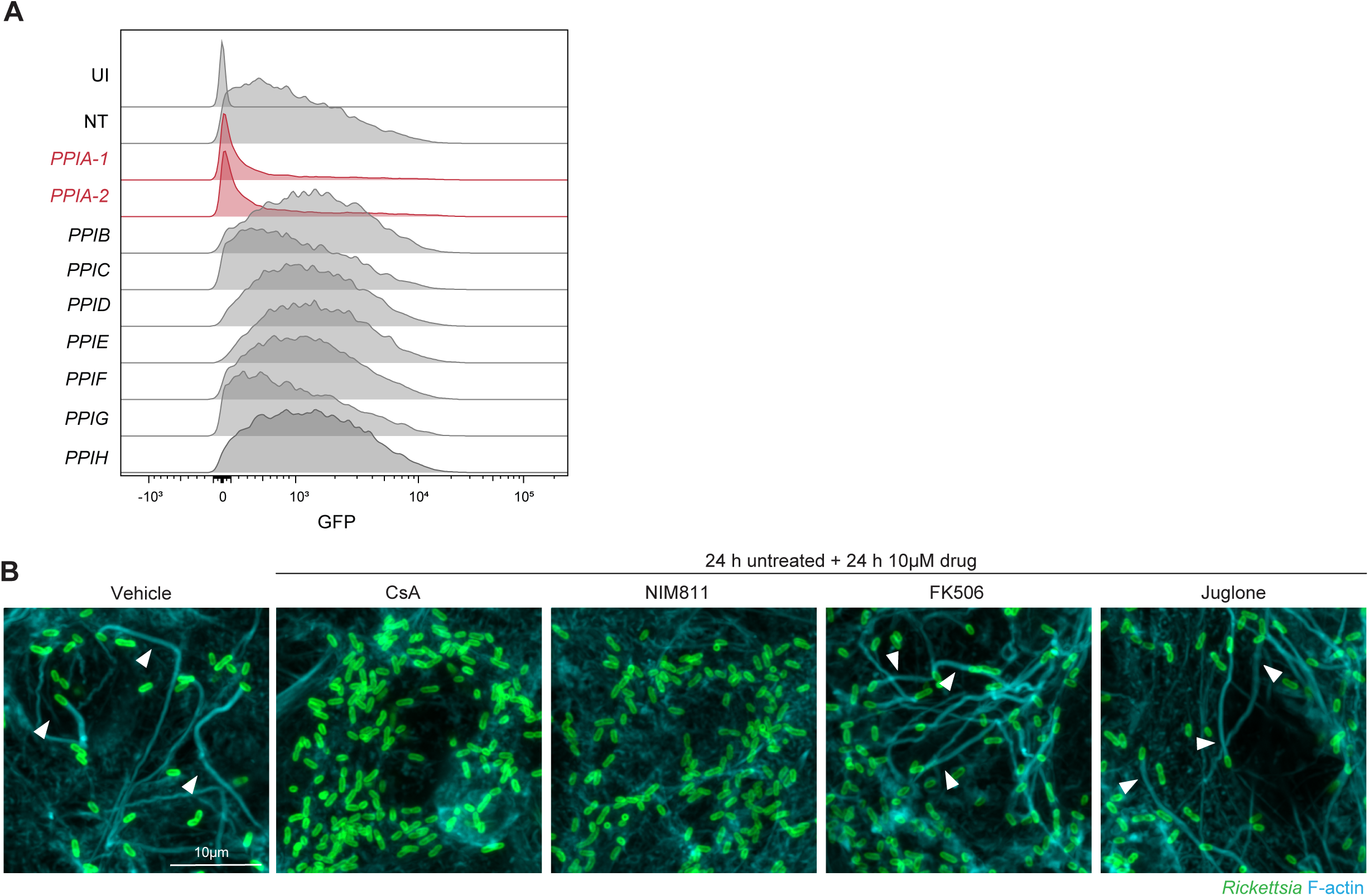
PPIA-specific dependence in *R. parkeri* infection and actin-based motility. (**A**): Flow cytometry curves of indicated knockout A549 lines infected with *Rp*AausFP1 for 48 hpi. *PPIA*-1 and *PPIA*-2 are two independent guides targeting *PPIA*. NT, non-target. Curves are representative of 3 independent infections. (**B**): Actin tails at 48 hpi in cells treated with 10μM cyclosporin A (CsA), NIM811, FK506 (FKBP inhibitor), or juglone (parvulin inhibitor) for 24 h starting at 24 hpi. Images are representative of n=3 biological replicates infected with WT *R. parkeri*. Arrowheads highlight bacteria with actin tails. (**A**): Comparison of Sca2 passenger domain structure to a canonical autotransporter (pertactin). Note the lack of β-stranded-regions in Sca2. Sca2 is colored according to Figure 4A layout. (**B**): Conservation of Gly-Pro containing repeats across selected *Rickettsia* spp. representative of different rickettsial clades. AG, ancestral group. TRG, transitional group. TG, typhus group. SFG, spotted fever group. Sca2 sequences were identified by protein BLAST with *R. parkeri* Sca2 and aligned with MAFFT. (**C**): Predicted structure of the PPIA interaction with Sca2(1087-1544) as the input, colored by pLDDT. PPIA is surface shaded. (**D**): Close-up view of PPIA-CRD interaction at second Gly-Pro site. (**E**): Atomic interactions at the predicted binding site between Sca2 Gly-Pro 1284-1285 and PPIA. Right: binding site of PPIA with HIV-1 capsid protein (PDB: 1AK4) Hydrogen bonds are indicated as blue dashed lines.

**Figure S4.**
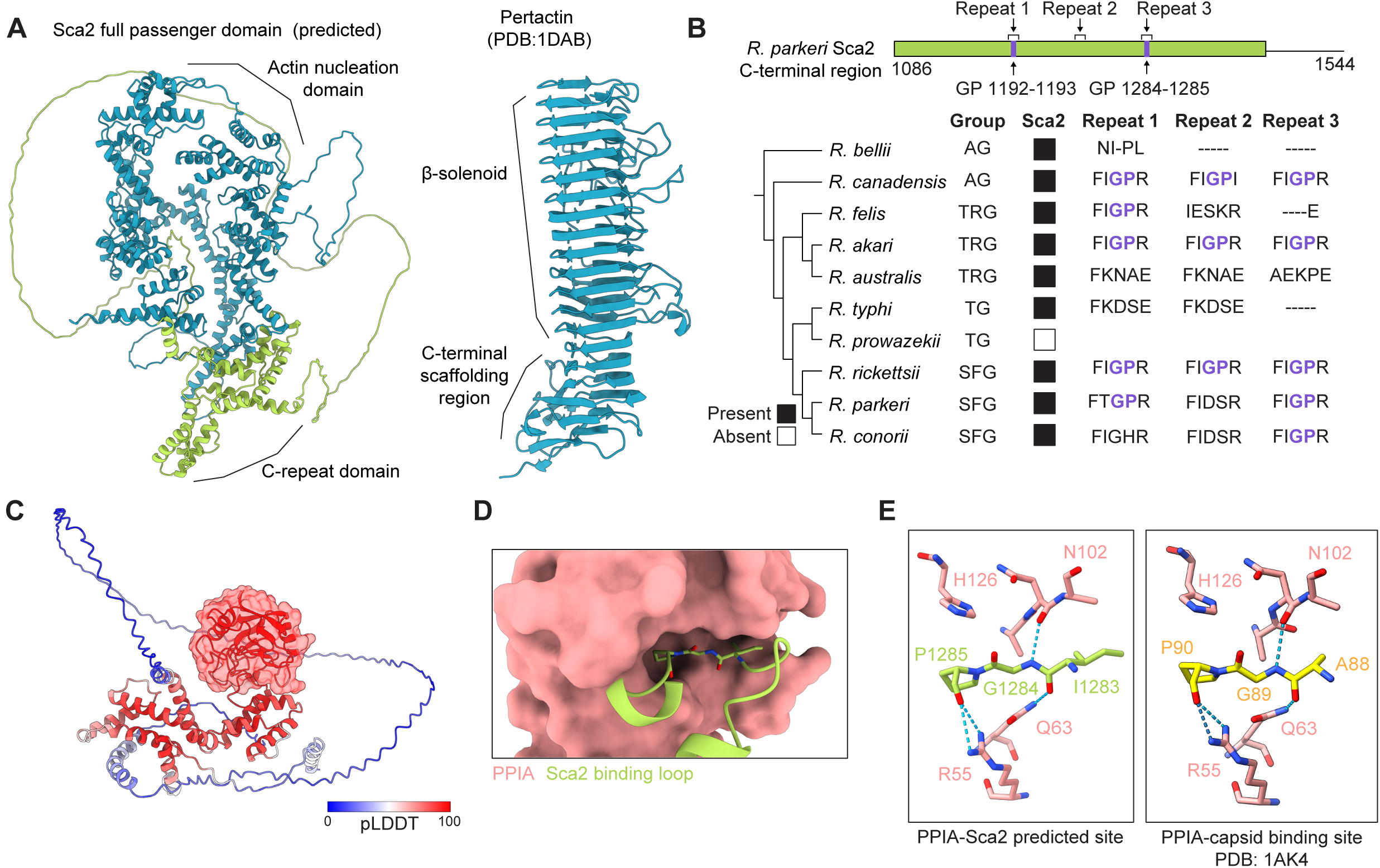
Sequence and structural elements of the Sca2-PPIA interaction. (**A**): Comparison of Sca2 passenger domain structure to a canonical autotransporter (pertactin). Note the lack of β-stranded-regions in Sca2. Sca2 is colored according to Figure 4A layout. (**B**): Conservation of Gly-Pro containing repeats across selected *Rickettsia* spp. representative of different rickettsial clades. AG, ancestral group. TRG, transitional group. TG, typhus group. SFG, spotted fever group. Sca2 sequences were identified by protein BLAST with *R. parkeri* Sca2 and aligned with MAFFT. (**C**): Predicted structure of the PPIA interaction with Sca2(1087-1544) as the input, colored by pLDDT. PPIA is surface shaded. (**D**): Close-up view of PPIA-CRD interaction at second Gly-Pro site. (**E**): Atomic interactions at the predicted binding site between Sca2 Gly-Pro 1284-1285 and PPIA. Right: binding site of PPIA with HIV-1 capsid protein (PDB: 1AK4) Hydrogen bonds are indicated as blue dashed lines.

**Figure S5.**
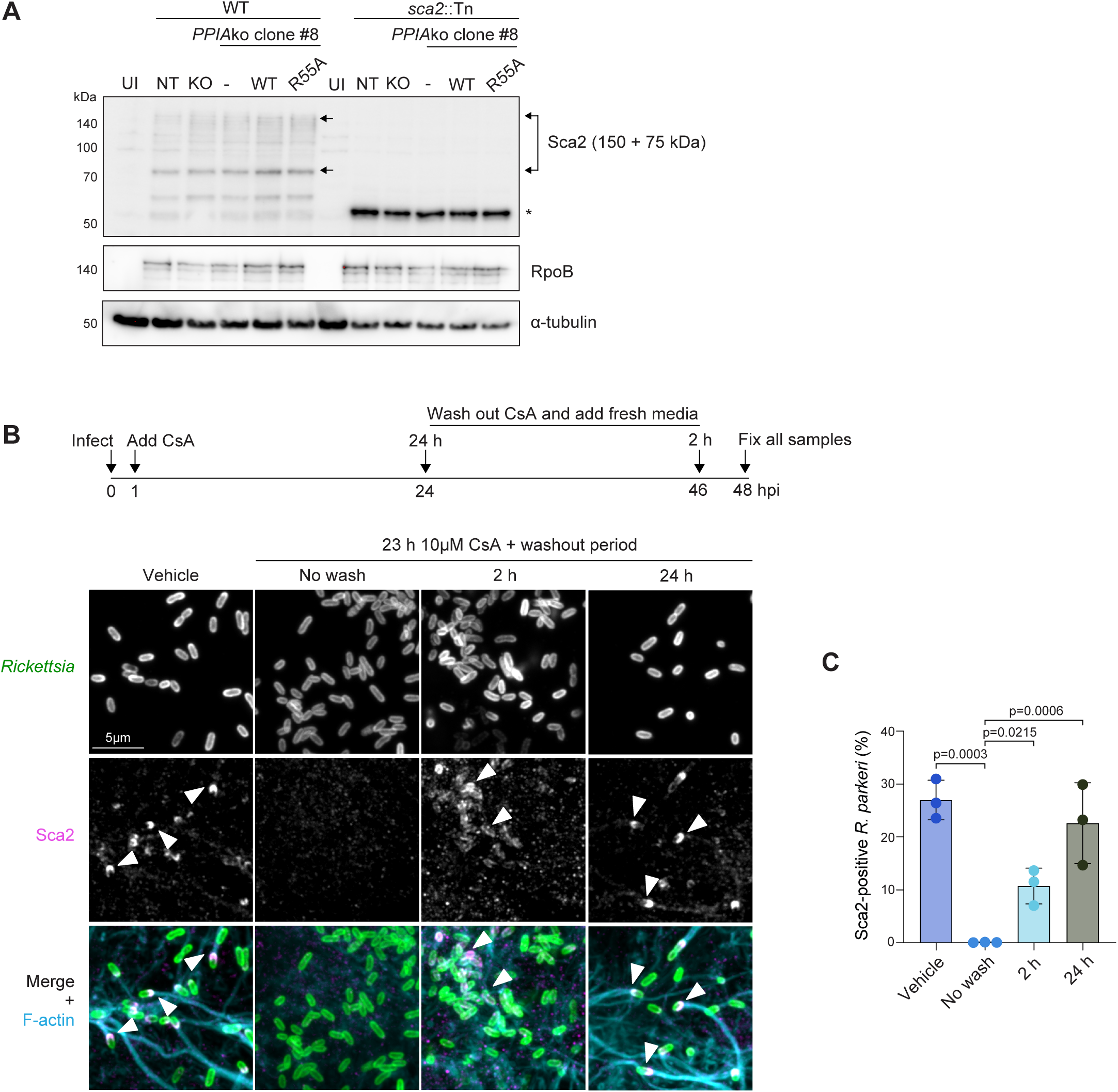
Additional characterization of Sca2 dependencies on PPIA. (**A**): Sca2 levels at 48 hpi with WT or *sca2*::Tn *R. parkeri* in NT or *PPIA*ko A549 host cells. Sca2 is known to blot as a 150kDa and 75kDa band (indicated with arrows). *: truncation product of Sca2 made by *sca2*::Tn bacteria. RpoB and tubulin are loading controls for *R. parkeri* and host input, respectively. UI: uninfected cells. Blot is representative of two independent experiments. (**B**): Sca2 surface exposure in WT A549 cells infected with WT *R. parkeri* treated with 10μM CsA for 24 h and washed out for the indicated time. Arrowheads highlight bacteria with Sca2 staining. (**C**): Quantification of data from Panel B. Means and SD from n=3 biological replicates (>2000 bacteria counted per replicate) are plotted. Means were used to calculate p-values (one-way ANOVA with Holm-Šídák multiple comparisons test). Images are representative of n=3 biological replicates.

**Figure S6.**
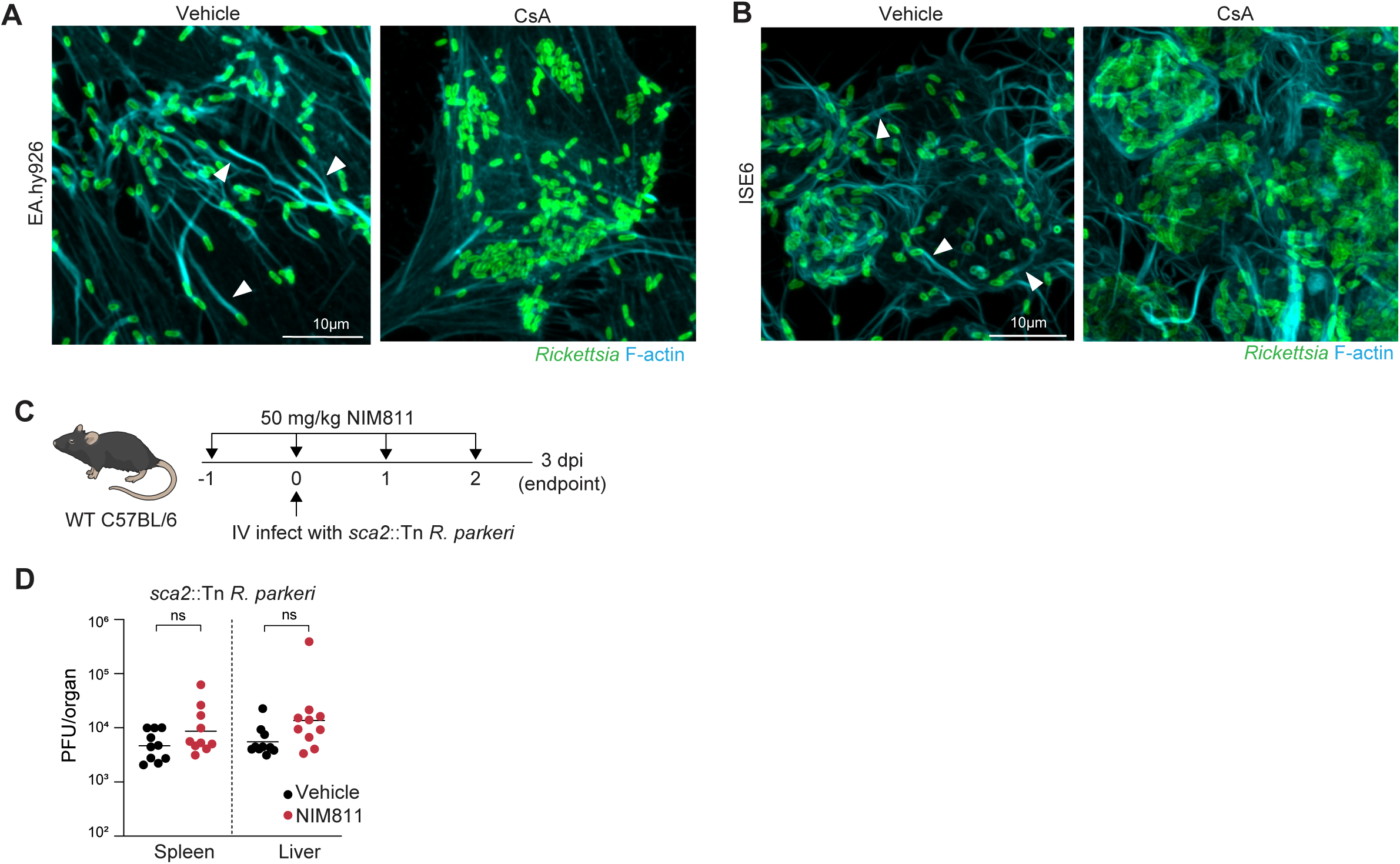
Additional characterization of PPIA in *R. parkeri* infection across cellular and *in vivo* contexts. (**A**): Actin tails at 48 hpi in EA.hy926 cells treated with vehicle or 10 μM CsA starting at 24 hpi. (**B**): Actin tails at 48 hpi in ISE6 cells treated with vehicle (DMSO) or 5 μM CsA starting at 1 hpi. (**C**): Schematic of *sca2*::Tn mouse infection experiment. (**D**): *sca2*::Tn *R. parkeri* plaque-forming units (PFU) in spleen (left) and livers (right) from infected mice. Each point represents one mouse. Data were aggregated from two independent experiments and analyzed with Mann-Whitney U tests. All images are representative of n=3 biological replicates infected with WT *R. parkeri*. Arrowheads highlight bacteria with actin tails.

